# Time or distance encoding by hippocampal neurons with heterogenous ramping rates

**DOI:** 10.1101/2023.03.12.532295

**Authors:** Raphael Heldman, Dongyan Pang, Xiaoliang Zhao, Brett Mensh, Yingxue Wang

## Abstract

To navigate their environments effectively, animals frequently integrate distance or time information to seek food and avoid threats. This integration process is thought to engage hippocampal neurons that fire at specific distances or times. Using virtual-reality environments, we uncovered two previously unknown functional subpopulations of CA1 pyramidal neurons that encode distance or time through a novel two-phase coding mechanism. The first subpopulation exhibits a collective increase in activity that peaks at similar times, marking the onset of integration; subsequently, individual neurons gradually diverge in their firing rates due to heterogeneous decay rates, enabling time encoding. In contrast, the second subpopulation initially decreases its activity before gradually ramping up. Closed-loop optogenetic experiments revealed that inactivating somatostatin-positive (SST) interneurons disrupts the first subpopulation, behaviorally impairing integration accuracy, while inactivating parvalbumin-positive (PV) interneurons disrupts the second subpopulation, impairing behavior during integration initiation. These findings support the conclusion that SST interneurons establish an integration window, while PV interneurons generate a reset to reinitiate integration. This study elucidates parallel neural circuits that facilitate distinct aspects of distance or time integration, offering new insights into the computations underlying navigation and memory encoding.

## INTRODUCTION

When navigating a dimly lit path at night, we often rely on our sense of distance traveled to determine our location. This process, known as path integration, involves the hippocampus integrating self-motion information into distance relative to recognizable landmarks, such as a distinctive tree or a notable building^1–4^.

In well-lit environments rich with cues, some hippocampal pyramidal neurons, called place cells, preferentially activate at specific locations^5^. Yet, even in dimly light environments or when environmental cues are sparse or absent, neurons can still activate at particular distances or time points relative to a landmark. These distance- or time-specific activity patterns are largely driven by internal computation rather than external cues, forming what are known as internally generated firing fields (IGFs)^6–11^. Collectively, neurons with IGFs activate in sequence to produce an internally generated sequence (IGS), which encodes moment-to-moment distance or time to aid path integration.

Recent evidence has begun to illuminate the circuit-level mechanisms supporting the activity of place cells and IGF neurons, with local inhibitory interneurons playing a key role. Different interneuron subtypes exhibit distinct temporal dynamics and target specific subcellular compartments of pyramidal neurons^12^. For example, somatostatin-positive (SST) interneurons primarily inhibit dendrites, whereas parvalbumin-positive (PV) interneurons preferentially inhibit the perisomatic region^12^. This compartment-specific inhibition regulates synaptic integration and spike output^13–16^, thereby shaping the spatial and temporal firing patterns of place cells^17–19^. These mechanisms likely apply to neurons with IGFs, which arise when environmental cues are absent.

However, only a subset of active pyramidal neurons exhibit IGFs, raising a critical question: What are the broader population dynamics during path integration, and how do inhibitory interneurons contribute?

In this study, we uncovered a novel neural code in the hippocampal CA1 area, featuring two previously unknown functional subpopulations of neurons. These populations track distance or time by first responding synchronously at integration onset, then diverging through ramping activity at different rates. By examining local inhibitory interneurons, specifically SST and PV interneurons^17–19^, we found that they played distinct roles in shaping these activity patterns and influencing the accuracy of path integration. Thus, our study reveals two parallel neural circuits within CA1 that coordinate to facilitate distance or time integration, offering new insights into the circuit-level mechanisms behind navigation and memory encoding.

## RESULTS

### A behavioral task that requires distance integration generates IGSs

We developed virtual reality (VR) setups (Figure 1A) that enable precise control over when and where visual stimuli are presented to a head-fixed mouse on a treadmill. Using these setups, we designed a behavioral task that requires animals to accumulate distance — referred to as a path integration (PI) task. Each trial starts with a brief visual cue, after which the animal must run 180 cm while the VR displays a constant grey background, creating a cue-constant segment. To receive a water reward, the animal has to lick a designated port within a reward zone; failure to do so results in no reward (Figure 1B left; Methods).

**Figure 1.**
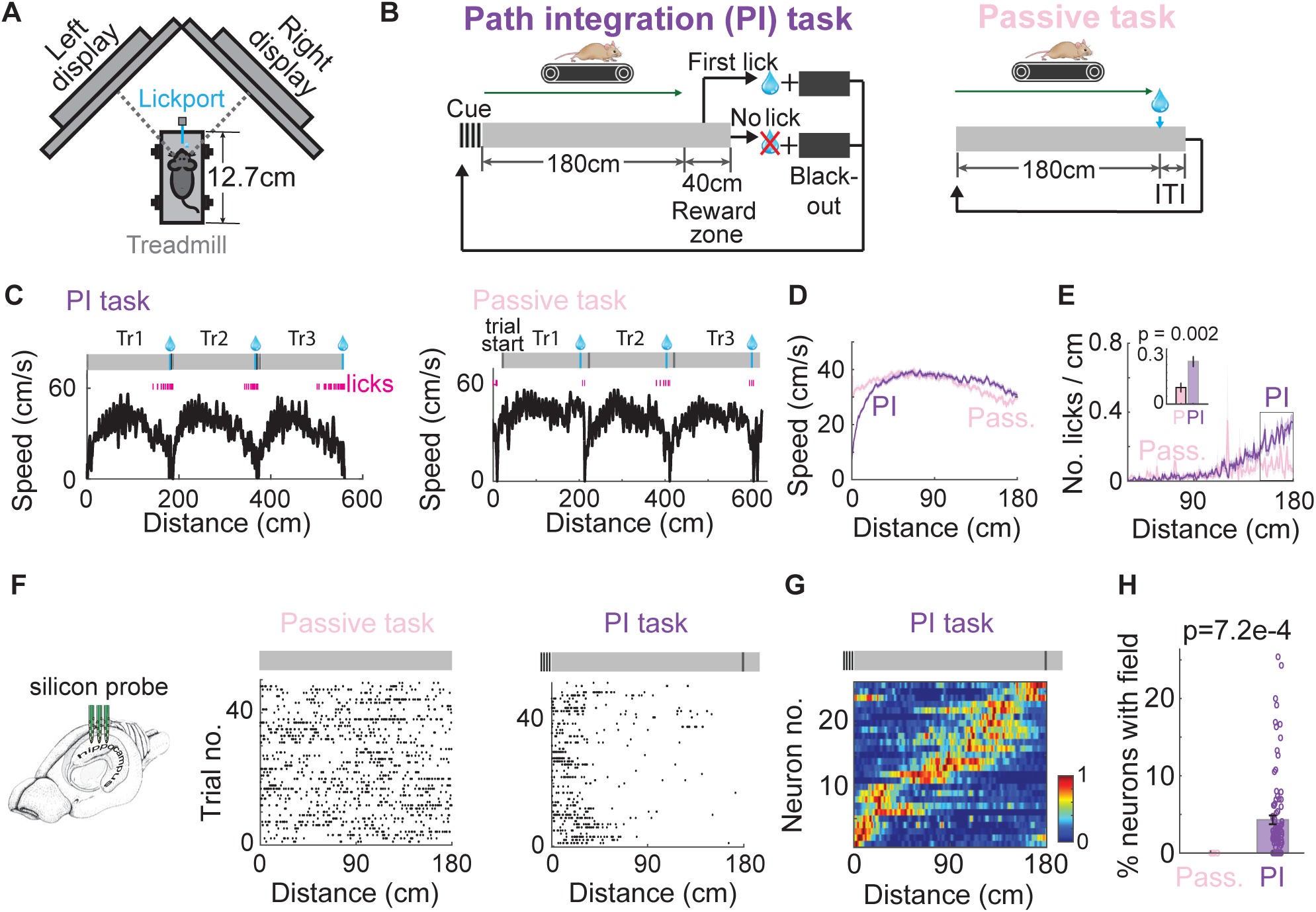
A path integration task that expresses internally generated fields (IGFs) (A) Schematic of the virtual reality setup: The mouse is head-fixed, with the eyes aligned with the two monitors. (B) Task schematics for PI (left) and passive (right) tasks (Methods). (C) Example speed traces (black) and licking activity (magenta) plotted against distance during three consecutive trials for mice performing the PI task (left) or passive task (right). TrN: Trial N. (D-E) Averaged speed traces (D) and lick histograms (E) across mice trained in either the passive (3 animals, 8 recordings, in pink) or PI task (4 animals, 10 recordings, in purple). The shaded area represents SEM (Same for other figures if not specifically stated). The inset bar plot compares the number of licks/cm within 150-180 cm between the two tasks. (F) Schematic of recordings with a silicon probe in the hippocampus (left). Spike rasters for example pyramidal neurons recorded in the passive (middle, no firing field) and PI tasks (right, with a field), plotted over distance within the cue-constant segment. (G) An example IGS within one session, plotting over distance within the cue-constant segment. Each row shows one neuron’s normalized firing rate change averaged over trials. (H) The percentage of neurons with fields per recording in the passive (3 animals, 8 recordings) and PI tasks (28 animals, 102 recordings). Each circle represents one recording.

Trained animals displayed stereotypical running speed profiles and licking patterns across trials (Figures 1C left, 1D and 1E), and accurate performance depended on an intact hippocampus (Figures S1A-S1D). Animals initiated licking as they approached the end of the cue-constant segment (Figure 1E, mean number of licks/cm from 100 to 180 cm: 0.17±0.022. Mean±SEM and rank-sum test are used if not specifically stated), implying that they track the distance to anticipate the reward.

Extracellular recordings in CA1 using silicon probes revealed that a subset of pyramidal neurons fired at specific distances during the cue-constant segment, expressing internally generated firing fields (IGFs) (Figures 1F right and 1H, 4.30±0.58% neurons; Methods). These neurons collectively form an IGS that encodes the moment-to-moment running distance (Figure 1G).

To verify whether IGS expression requires distance integration, we compared the PI task with a passive task. In the passive task, animals run for 180 cm while viewing the grey screens, and a water reward is automatically delivered at the end without requiring licking (Figure 1B right, Methods). This task does not require animals to integrate over distance and replicates previously reported non-memory tasks^6,20^. Animals ran at a similar mean running speed to the PI task (Figures 1C right and 1D, Methods, Passive: 34.41±1.80 cm/s, PI: 34.73±1.23 cm/s, p=0.63, rank-sum test). However, in the passive task, there was a significant decrease in predictive licks before reward delivery (Figure 1E, mean number of licks per cm from 100 to 180 cm, Passive: 0.083±0.025, PI: 0.17±0.022, p=0.012.).

In the passive task, CA1 pyramidal neurons exhibited significantly lower trial-by-trial activity correlation (Figure S1E, mean activity correlation over trials, Passive: 0.006±7.9e-04 (536 neurons), PI: 0.033±5.2e-4 (7828 neurons), p=3.6e-94). These neurons fired throughout the cue-constant segment (Figure 1F middle), and the IGF was absent (Figure 1H). Thus, IGSs preferentially occur during the task requiring distance integration.

### The initiation of an IGS aligns with self-initiated onset of running

In the PI task, the run onset potentially marked the starting point of integration in the PI task (Figure 2B). Since the interval between the initial visual cue and run onset often did not align — on average, there was a delay of 1.15 seconds (Figures 2A-2D, Methods) — animals likely self-initiated their running rather than merely responding to the cue.

**Figure 2.**
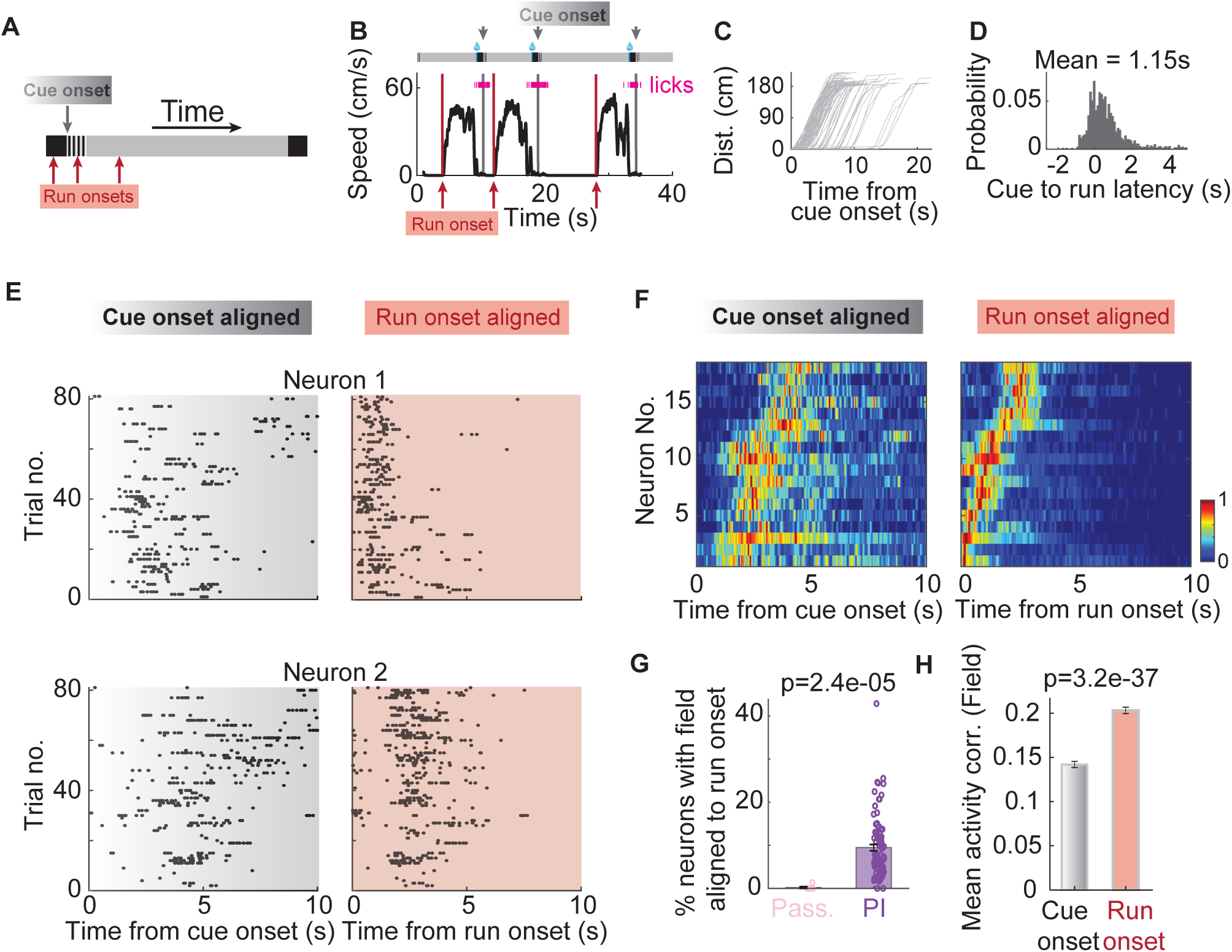
Internally generated sequence (IGS) starts at run onset in the PI task. (A) The relationship between cue onset and run onset can be resolved when examining each trial in time instead of distance. Grey arrow: cue onset; red arrows: possible run onset times. (B) Example speed traces and licking behavior plotted against time unfolded across 3 consecutive trials, where run onset does not align with cue onset. Run onsets: red arrows. Cue onsets: grey arrows. (C) Distance traveled as a function of time from cue onset for all trials within one recording session. Each grey line corresponds to one trial. (D) Probability distribution of the time from cue onset to run onset (28 animals, 102 recordings). (E-F) Spiking activity of two example pyramidal neurons (E), and an example IGS (F) from a single session (the same session as in (C)), aligned with cue onset (left) vs. run onset (right). (G) The percentage of pyramidal neurons with IGFs per recording in the passive and PI tasks after aligning with run onset. (H) Mean activity correlation of neurons with IGFs when aligned with cue vs. run onset (650 neurons, cue onset: 0.14±0.0035, run onset: 0.20±0.0037).

Aligning neuronal activity with run onset in each trial revealed consistent firing at specific time points (IGFs). In contrast, these IGFs were less evident when aligning the same activity with cue onset (Figure 2E). Under run-onset alignment, more neurons exhibited IGFs (Figure 2G) and showed higher trial-by-trial activity correlation (Figure 2H). We also observed increased temporal information (cue onset: 0.29±0.0077bits/s, run onset: 0.34±0.0091bits/s, p=3.0e-05), and more pronounced phase precession^6,21,22^ (Figures S1F and S1G). At the population level, IGSs became sharper with run-onset alignment (Figure 2F, population correlation: cue onset: 0.20±0.0086, run onset: 0.24±0.0076, p = 1.9e-04).

In contrast, IGFs remained absent in the passive task even when aligning activity with run onset (Figure 2G, PI: 9.46±0.72% neurons, Passive: 0.19±0.19% neurons; Methods). This result suggests that engaging in distance integration is necessary for generating IGFs. However, it remains ambiguous whether animals were integrating over distance or time (Figures 3C left and S2B left), as they maintained consistent running speed curves across trials.

**Figure 3.**
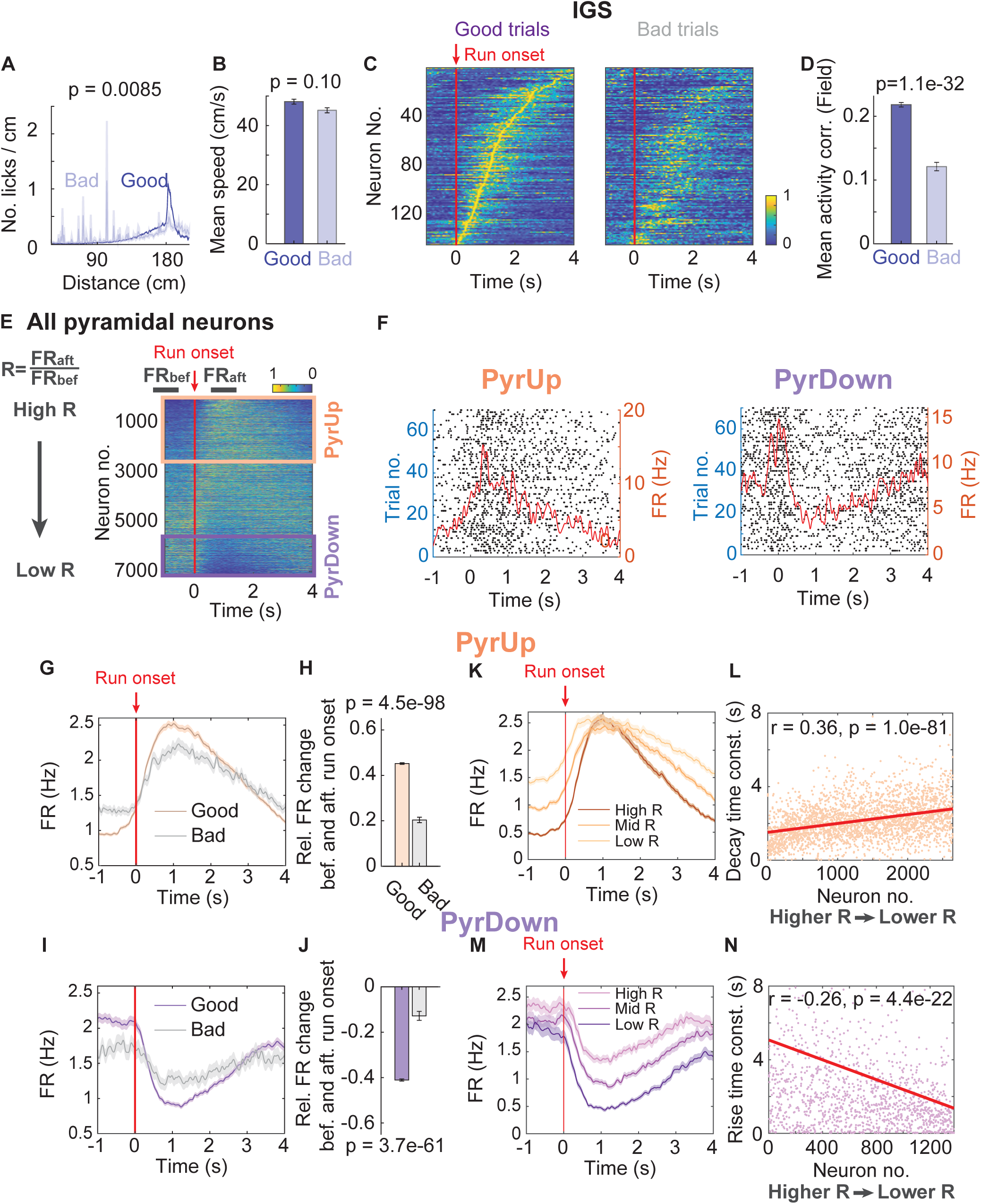
CA1 pyramidal neurons display distinct responses around the start of integration. (A-B) Lick histogram (reported p-value is for mean number of licks/cm between 30-100 cm) (A) and mean running speed (B), broken down by good (blue) and bad trials (light blue). (C) IGS formed by neurons with IGFs from all the recordings with at least 15 bad trials (15 animals, 22 recordings), averaged across good (left) and bad (right) trials. (D) Mean trial-by-trial activity correlation of neurons with IGFs, broken down by good (blue) and bad trials (light blue). Good trials: 0.22±0.0035, bad trials: 0.12±0.0066. (E) After aligning with run onset, normalized firing rate heatmaps of all pyramidal neurons ordered by their firing rate ratio R = FR_aft_/FR_bef_ (FR_bef_: the averaged firing rate (FR) before run onset (−1.5 to −0.5s), FR_aft_: FR after run onset (0.5 to 1.5s), see black bars on the top, 0s corresponds to run onset). Recordings with at least 15 good trials are included. PyrUp (2645 neurons) and PyrDown (1377 neurons) neurons are denoted by orange and purple boxes, respectively. (F) Spike rasters for example PyrUp (left) and PyrDown (right) neurons. The averaged firing rate over time is superimposed (red line). (G) The firing rate profile averaged across all PyrUp neurons broken down into good and bad trials. PyrUp neurons are defined as neurons with R > 3/2, the presence of PyrUp was not sensitive to the threshold value. The shaded area shows the SEM. (H) Relative firing rate change before and after run onset (FRbef – FRaft)/(FRbef + FRaft) for PyrUp neurons. Good trials: 0.45±0.0036, bad trials: 0.20±0.012. (I-J) Same as in (G-H), for all recorded PyrDown neurons. Good trials: −0.41±0.0048, bad trials: −0.13±0.020. PyrDown neurons are defined as neurons with R < 2/3, the presence of PyrDown was not sensitive to the threshold value. (K) Dividing PyrUp neurons into three equal-sized groups based on their R values: high R, mid R, and low R (dark to light lines). Comparing the average firing rate profile among these groups. Peak activity time of the high R vs low R groups: p = 0.47. (L) The decay time constant for each PyrUp neuron’s firing rate profile following its peak is shown in orange dots (Methods). Neurons are arranged in the order of descending R values. The linear regression line (red) illustrates the change of decay time constants across neurons. (M-N) Same as (K-L), for all the PyrDown neurons.

Moreover, the alignment with run onset enhanced trial-by-trial activity correlations not only in neurons with IGFs but across all recorded CA1 pyramidal cells (Figures S1H and S1I). This effect is unlikely due to a switch from a non-theta to theta brain state when animals transitioned from reward collection at the end of one trial to starting the next run, because the average pause time was brief (pause mean time = 1.88s, Figures S1J and S1K).

In summary, run onset aligns the neuronal activity involved in path integration, likely marking the initiation of the neural computation for distance or time estimation.

### Accurate integration is correlated with the expression of IGSs

To investigate how IGS expression relates to task performance, we categorized each trial as either “good” or “bad”. Good trials met the following criteria: the animals came to a complete stop before trial run onset, ran largely uninterrupted, and successfully obtained the reward (Methods). These trials constituted 77.90±2.88% of all trials and had a 100% reward rate. In contrast, bad trials failed at least one criterion and had a lower reward rate of 86.34±4.36%.

In bad trials, animals showed more licking in the early part of running, despite maintaining a mean running speed similar to that in good trials (Figures 3A and 3B, mean licks/cm before 100 cm distance: good trials: 0.011±0.0024, bad trials: 0.027±0.0063; mean licks/20ms between 0 to 1s time: good trials: 0.012±0.0021, bad trials: 0.041±0.0049, p=1.36e-8; mean speed: good trials: 48.12±0.84cm/s, bad trials: 45.20±0.84cm/s). This behavior implied an inaccurate estimation of distance or time.

Correspondingly, in bad trials, neurons with IGFs showed reduced trial-by-trial activity correlation (Figure 3D), and the IGSs were impaired (Figures 3C and S2A). The impairment persisted when IGSs were plotted against distance instead of time (Figures S2B and S2C). Together, these findings indicate that the expression of IGSs is closely linked to accurate distance or time estimation, as reflected in both neuronal and behavioral measures.

### Subpopulations of pyramidal neurons display distinct responses around run onset

In addition to IGSs, CA1 pyramidal neurons generally exhibited improved trial-by-trial activity correlation when aligned to run onset (Figures S2H). To visualize how overall the pyramidal neurons dynamically changed their activity over time or distance from run onset, we sorted all recorded neurons (7114 neurons, 28 animals, including neurons with IGFs (650 neurons)) according to the ratio (R) of their mean firing rates after versus before run onset (Figure 3E and S2D).

This approach revealed two distinct functional subpopulations of pyramidal neurons. “PyrUp” neurons (37.2% of the neurons) rapidly increased their activity at run onset and then gradually ramped down as the animal approached the reward (Figures 3F left, 3G, and 3H), whereas “PyrDown” neurons (19.4%) initially reduced their activity but then ramped up toward reward (Figures 3F right, 3I, and 3J). Neurons that did not fit either pattern were labeled “PyrOther” (Figure S2E). The results were similar when we restricted our analysis to neurons whose R values significantly deviated from shuffled data (Figures S2G-S2I; Methods).

Both PyrUp and PyrDown subpopulations displayed more pronounced firing rate changes around run onset in good trials compared to bad trials (Figures 3G-3J), despite similar overall firing rates across trial types (measured from −1 to 4 seconds, PyrUp: good trials: 1.76±0.032Hz, bad trials: 1.72±0.061Hz, p=0.19, PyrDown: good trials: 1.49±0.042Hz, bad trials: 1.50±0.087Hz, p=0.27). That is, their response strength correlated with the accuracy of distance or time estimation. Moreover, PyrUp and PyrDown did not differ significantly in their relative position within the CA1 pyramidal layer^23^ (Figure S2F), implying that each subpopulation likely includes both genetically distinct deep and superficial CA1 neurons^23–27^.

To assess whether these subpopulations encode time, we applied Bayesian decoding^28^ to their single-trial population activity. Focusing on PyrUp neurons without IGFs, we confirmed that they effectively encode time passage (Figures S2J-S2M). However, unlike the sequential activation of neurons with IGFs at specific time points, PyrUp neurons exhibited a collective peak at similar times shortly after run onset (Figure 3K, peak time: mean±SEM = 1.24 ± 0.02 s). Their activity then decayed at heterogeneous, neuron-specific rates. Neurons with slower decay rates tended to have lower R values (Figures 3K and 3L) and higher baseline firing before run onset (r = 0.26, p = 1.18e-41). Based on these observations, we hypothesize that PyrUp neurons encode time through a two-phase strategy: (1) an integration “start” phase marked by the synchronized increase in activity that peaks around similar times across the population. (2) a time-encoding phase characterized by the gradual divergence in individual neurons’ firing rates, with greater divergence corresponding to a longer elapsed time (Figure 3K).

A similar coding mechanism was seen in PyrDown neurons, where those that increased their activity with faster rates toward reward tended to have lower R values (Figures 3M and 3N).

In summary, we uncovered two novel functional subpopulations, PyrUp and PyrDown neurons. These neurons exhibit a synchronized start response followed by activity increase or decrease with heterogeneous rates. This synchronized “start” provides a reference point for subsequent time or distance encoding. This coding mechanism is distinct from the sequential, point-by-point encoding through neurons with IGFs.

### PyrUp and PyrDown responses occur specifically at the start of integration

We further asked whether the PyrUp and PyrDown responses observed around the trial-start run onset (TRO) apply to any run onset. At the spontaneous run onset (SRO) occasionally occurring in the middle of a trial (19 animals, 32 sessions, 2902 neurons; Methods), both responses were significantly attenuated compared to TRO, despite similar mean firing rates (Figures S3A and S3B, PyrUp: at SRO: 2.12±0.06Hz, at TRO: 2.11±0.06Hz, PyrDown: at SRO: 1.70±0.088Hz, at TRO: 1.72±0.094Hz) and matching running speed (Figure S3C, 13 animals, 21 sessions, 1731 neurons; Methods). These results suggest that PyrUp and PyrDown responses are context-dependent and specifically tied to run onset when the distance or time integration starts.

To verify that the PyrUp and PyrDown responses were not merely a result of locomotion, we designed an immobile task where the animal remained stationary^29^ and had to estimate a 4-second time period while viewing grey screens (Figure S3D; Methods). Even in this immobile state, we observed IGSs (5 animals, 15 sessions, 748 neurons, activity aligning with the last lick of the previous trial, Figure S3E, 6.72±1.56% neurons) and high percentages of neurons with PyrUp and PyrDown responses (Figures S3D-S3J, PyrUp-Imm: 50.71±4.18% or “PyrDown-Imm: 26.18±2.60%).

Altogether, these results suggest that PyrUp and PyrDown responses occur during tasks requiring time or distance integration across distinct behavioral states, including locomotion and immobility.

### Somatostatin-positive interneurons display slow ramping activity around run onset

To reveal the circuit-level mechanisms of the PyrUp and PyrDown responses, we first examined local somatostatin-positive (SST) inhibitory interneurons. These interneurons target the dendrites of pyramidal neurons and coordinate the interaction between inputs from CA3 and the entorhinal cortex^12,17,18^.

To identify SST interneurons, we used optogenetic tagging by transfecting the CA1 SST interneurons with channelrhodopsin-2 (ChR2) in SST-IRES-Cre animals (Figures 4D and S4B, 3 animals). Meanwhile, we classified putative inhibitory neurons^14^ by their spike properties and relative locations to the pyramidal layer center^12^ (Figures 4A and S4A, Methods), and identified a cluster that best overlapped with the tagged neurons (12/24 tagged neurons). Neurons in this cluster exhibited characteristics consistent with SST-expressing oriens-lacunosum moleculare (OLM) cells^12^ (Figures S4C and S4D).

**Figure 4.**
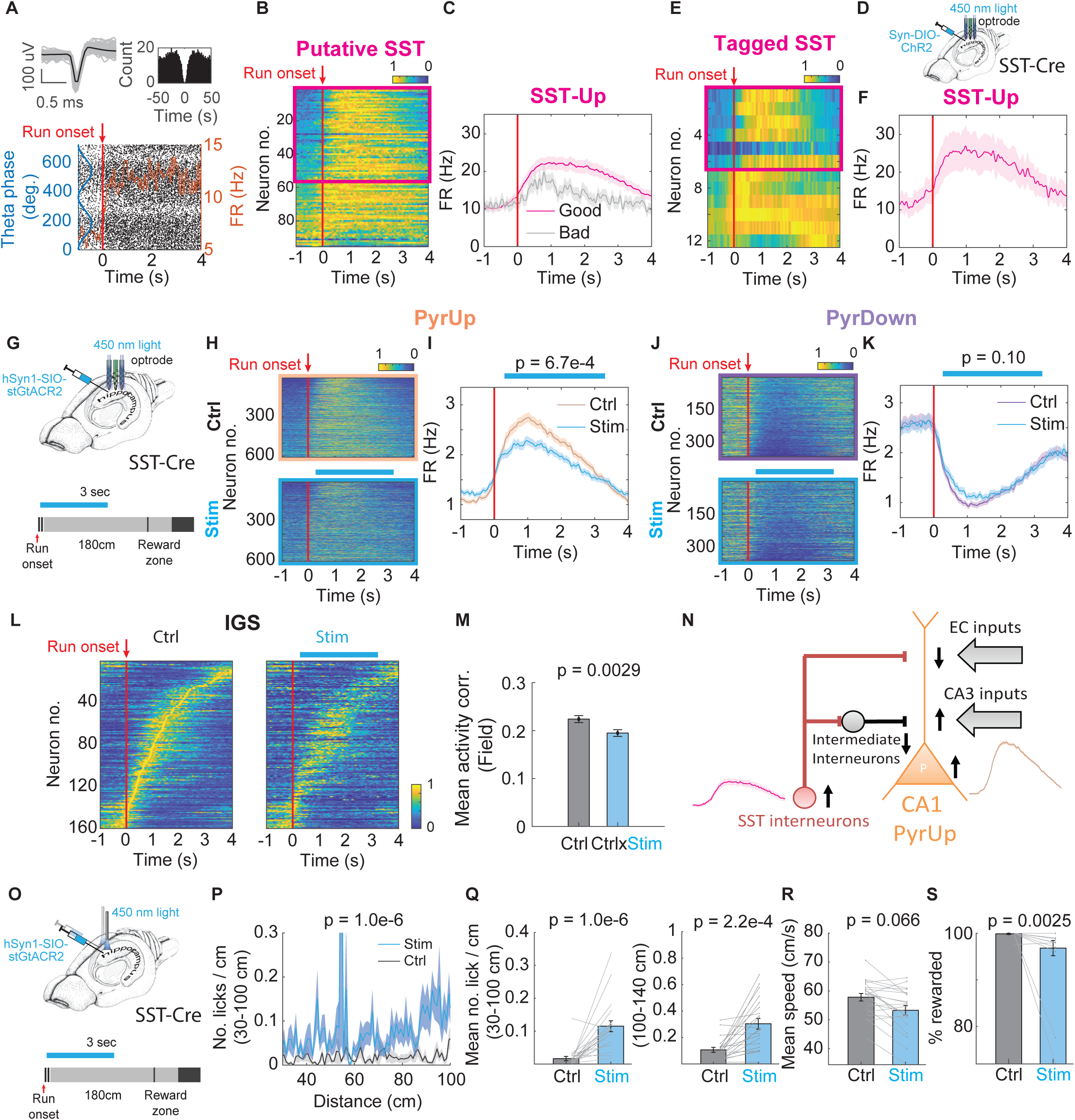
CA1 somatostatin-positive interneurons regulate PyrUp response, IGS, and distance integration. (A) Spike waveform (top-left) and auto-correlogram (top-right) of a putative SST interneuron. In top-left, single waveforms are shown in grey and the averaged waveform is shown in black. Raster plot (bottom) showing spike theta phases (black dots) and firing rate as a function of time (orange) for the same cell. (B) Normalized firing rate heatmaps of all the interneurons belonging to the putative SST interneuron cluster, ordered by their firing rate ratio FR_aft_/FR_bef_ around run onset. The magenta box denotes SST-Up neurons whose FR_aft_/FR_bef_ > 3/2. (C) Firing rate profile averaged across all SST-Up neurons broken down into good and bad trials. Relative FR change: good trials: 0.35±0.015, bad trials: 0.076±0.054, p=6.2e-06. (D) Schematic for the opto-tagging experiment. (E) Same as in (B), but for interneurons identified as SST-positive from the tagging experiments and at the same time belonging to the putative SST interneuron cluster (6/12 neurons with FR_aft_/FR_bef_ > 3/2). (F) Same as in (C), for tagged SST-Up neurons highlighted within the magenta box in (E). Bad trials are not shown because the number of bad trials is less than 15 trials in most of these recordings. (G) Experimental setup for inactivating SST-interneurons. Light stimulation is applied with an optrode for 3 seconds starting after run onset in a subset of trials (Methods). (H) Normalized firing rate heatmaps of all PyrUp neurons during control (top) and stimulation trials (bottom). Neurons are ordered by their firing rate ratio around run onset. The blue horizontal bar illustrates the estimated stimulation window. (I) Firing rate profile averaged across PyrUp neurons for control (orange) and stimulation trials (blue). The two-sample t-test compares control and stimulation trials, performed for the mean FR between 0.5-2s: control: 2.55±0.086Hz, stimulation: 2.15±0.077Hz, p=6.7e-04. (J) Same as in (H), for PyrDown neurons. (K) Same as in (I), for PyrDown neurons. mean FR between 0.5-2s: control: 1.05±0.056Hz, stimulation: 1.19±0.065Hz, p=0.10 (two-sample t-test). (L) IGS during control (left) and stimulation trials (right). (M) Mean activity correlation of neurons with IGFs, comparing control (grey) and stimulation trials (blue). Mean activity corr.: control: 0.22±0.0073, stimulation: 0.19±0.0071, p=0.0029. (N) A circuit diagram illustrates how SST inactivation potentially impacts PyrUp response. SST interneurons, by inhibiting intermediate interneurons, disinhibit the CA3 inputs to PyrUp neurons. (O) Experimental setup to assess the behavioral effects of SST inactivation. The same stimulation protocol as in (G) was applied to a different cohort of mice with optic fibers implanted bilaterally. (P) Lick histogram for the first 100 cm for control (grey) and stimulation (blue) trials (6 animals, 21 recordings). (Q-S) Mean number of licks/cm calculated based on (P) (Q left) (control: 0.017±0.0066, stimulation: 0.12±0.017), mean number of licks/cm between 100-140 cm (Q right) (control: 0.11±0.019, stimulation: 0.30±0.041), mean running speed (R) (control: 57.82±1.31cm/s, stimulation: 53.29±1.68cm/s) and percentage of rewarded trials (S) for control and stimulation trials (control: 99.88±0.12%, stimulation: 96.82±1.63%).

A majority of neurons in this cluster (55/95 neurons, Figures 4B and 4C), including half of the tagged cells (6/12 neurons, Figures 4E and 4F), increased their activity at run onset, which subsequently decayed as the reward was near in the PI task (SST-Up neurons). In bad trials, the SST-Up response was weakened (Figure 4C, mean FR: good trials: 17.52±1.42Hz, bad trials: 17.84±4.19Hz, p=0.95), suggesting its necessity for accurately estimating distance or time.

Previous evidence suggests that OLM interneurons directly inhibit entorhinal cortex inputs to CA1 pyramidal neurons while indirectly disinhibiting their CA3 inputs through other inhibitory interneurons^15,16,30^. Therefore, the period in which SST-Up neurons displayed elevated firing after run onset likely favors CA3 inputs.

### Inactivating SST interneurons after run onset impairs PyrUp response, IGSs, and task performance

To determine how SST activity influences PyrUp and PyrDown responses, we utilized soma-targeted anion-conducting channelrhodopsins (stGtACR2) to hyperpolarize SST interneurons^31^. We recorded the neuronal activity unilaterally from dorsal CA1 using an optrode (Figure 4G top) while performing closed-loop inactivation of SST interneurons for 3 seconds immediately after run onset (Figure 4G bottom, Methods). This manipulation effectively silenced some putative SST cells (Figures S4I and S4J).

Silencing SST interneurons significantly reduced the PyrUp response after run onset (Figures 4H and 4I, 6 animals, 24 recordings, 608/1721 neurons), but did not produce a clear effect on the PyrDown response (Figures 4J and 4K, 359/1721 neurons).

Neurons with IGFs showed a significant decrease in their trial-by-trial activity correlation without a change in overall firing rate (Figures 4L and 4M, 160/1721 neurons, mean FR: control: 1.99±0.11Hz, stimulation: 1.97±0.12Hz, p=0.44). These inactivation effects on the PyrUp response and IGSs resembled the corresponding results in bad trials (Figures 3C and 3G).

To assess behavioral consequences, we bilaterally silenced SST interneurons starting at run onset for 3 seconds in a separate cohort of animals (Figure 4O, Methods). In the stimulation sessions, animals showed a significant increase in early licks before 100 cm (Figures 4P and 4Q left, 6 animals), predictive licks between 100-140 cm (Figure 4Q right), and a decrease in rewarded trials (Figure 4S). However, mean running speed did not significantly change, indicating normal locomotion (Figure 4R). Control experiments ruled out that the effects were due to laser light alone (Figures S5F-S5J, S5P-S5T; Methods).

Overall, elevated SST activity after run onset is necessary for producing the PyrUp response and IGSs (Figure 4N). Reducing SST activity impairs the PyrUp response and IGSs, resulting in a disruption of distance or time integration.

### Inactivating SST interneurons close to reward does not affect PyrUp response or task performance

As SST activity declined near the reward (Figure 4N), CA3 inputs to PyrUp neurons may be gradually blocked. Thus, SST interneurons may establish a critical time window, spanning from run onset to near the reward, that facilitates the PyrUp response. Supporting this hypothesis, silencing SST interneurons for 3 seconds starting at 120 cm (Figure 5A) had minor effects on PyrUp neurons, PyrDown neurons (Figures 5B-5E, 6 animals, 18 recordings) and IGSs (Figures 5F and 5G, 161/1516 neurons, mean FR: control: 1.71±0.088Hz, stimulation: 1.71±0.099Hz, p=0.52).

**Figure 5.**
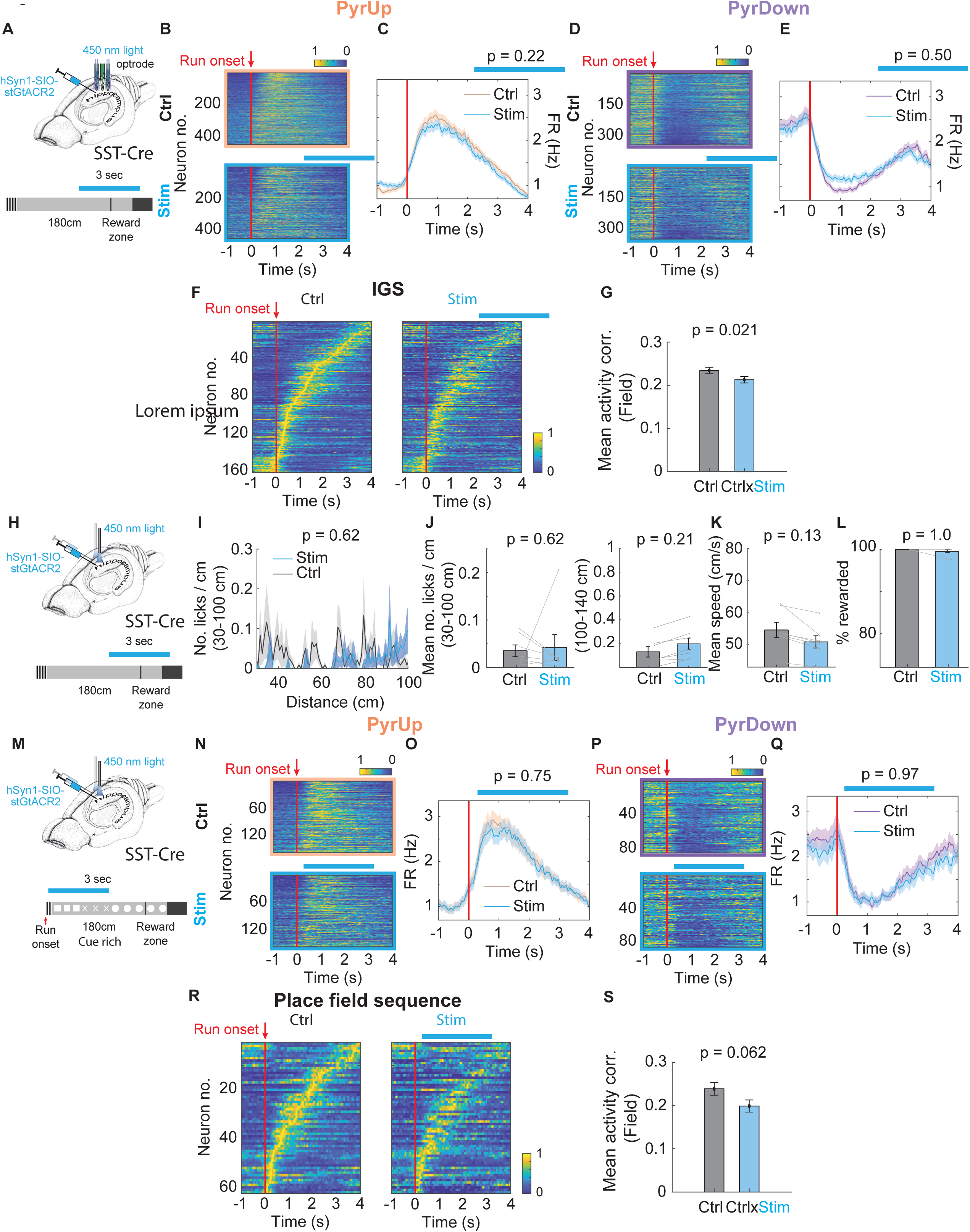
Inactivating CA1 somatostatin-positive interneurons close to the reward zone has minor impacts. (A) Experimental setup for inactivating SST interneurons. Light stimulation is applied with an optrode for 3 seconds starting at 120 cm in a subset of trials. (B-L) The same as Figures 4H-4M, 4O-4S, for SST inactivation after 120 cm as shown in (A). (B-E) PyrUp: 454/1516 neurons; mean FR between 2.5-4s: control: 1.22±0.058Hz, stimulation: 1.26±0.055Hz, p=0.22 (two-sample t-test); PyrDown: 348/1516 neurons; mean FR between 2.5-4s: control: 1.72±0.088Hz, stimulation: 1.63±0.082Hz, p=0.50 (two-sample t-test). (G) Mean activity corr.: control: 0.23±0.0074, stimulation: 0.21±0.0074, p=0.021. (I-L) including 4 animals, 7 recordings. The blue horizontal bar illustrates the estimated stimulation window. (M) Experimental setup for inactivating SST interneurons in a cue-rich environment. Visual cues are displayed on the screens (square, X, and circle, each occupying 60 cm of the 180 cm track). Light stimulation is applied with an optrode for 3 seconds starting after run onset in a subset of trials. (N-S) The same as Figures 4H-4M, for inactivating SST interneurons in a cue-rich environment as shown in (M). (N-Q) PyrUp: 158/422 neurons; mean FR between 0.5-2s: control: 2.63±0.18Hz, stimulation: 2.55±0.18Hz, p=0.75 (two-sample t-test); PyrDown: 86/422 neurons; mean FR between 0.5-2s: control: 1.10±0.093Hz, stimulation: 1.09±0.11Hz, p=0.97 (two-sample t-test). (S) Mean activity corr.: control: 0.24±0.015, stimulation: 0.20±0.014, p=0.062.

Similarly, bilaterally inactivating SST interneurons after 120 cm using the same cohort of animals as in Figure 4O did not alter animals’ behavior (Figures 5H-5L).

Therefore, SST interneurons are necessary for accurate distance or time integration, primarily within the time window from run onset to when animals approach the reward zone.

In our results, SST inactivation after run onset impaired IGSs. However, previous studies conducted in a cue-rich environment reported no clear disruption in place field sequences^18^. To reconcile this discrepancy, we introduced visual cues during the cue-constant segment of the PI task (Figure 5M) and repeated SST inactivation after run onset. Despite a shutdown of some putative SST cells, we observed no significant changes in neuronal sequences or the PyrUp response (Figures 5N-5S, 3 animals, 9 recordings). Our results suggest that the reduced PyrUp response mediated by SST inactivation is specific to internally generated dynamics, rather than response driven by external cues.

### Parvalbumin-positive cells display ramping activity around run onset

Because SST inactivation did not significantly affect the PyrDown neurons, we asked whether parvalbumin-positive (PV) interneurons, which primarily target the perisomatic region and regulate the spike timing and synchrony of pyramidal neurons^32–34^, may play a role in controlling the shutdown of PyrDown neurons at run onset.

To identify PV cells, we used optogenetic tagging by transfecting the CA1 PV interneurons with ChR2 in PV-Cre animals (Figures 6D and S4E; 7 animals). A large proportion of these tagged neurons overlapped with one putative inhibitory neuron cluster located primarily in the pyramidal layer (13/30 tagged neurons, Figures 6A and S4A). Neurons in this cluster exhibited characteristics consistent with PV basket cells (Figures S4F and S4G)^12^.

**Figure 6.**
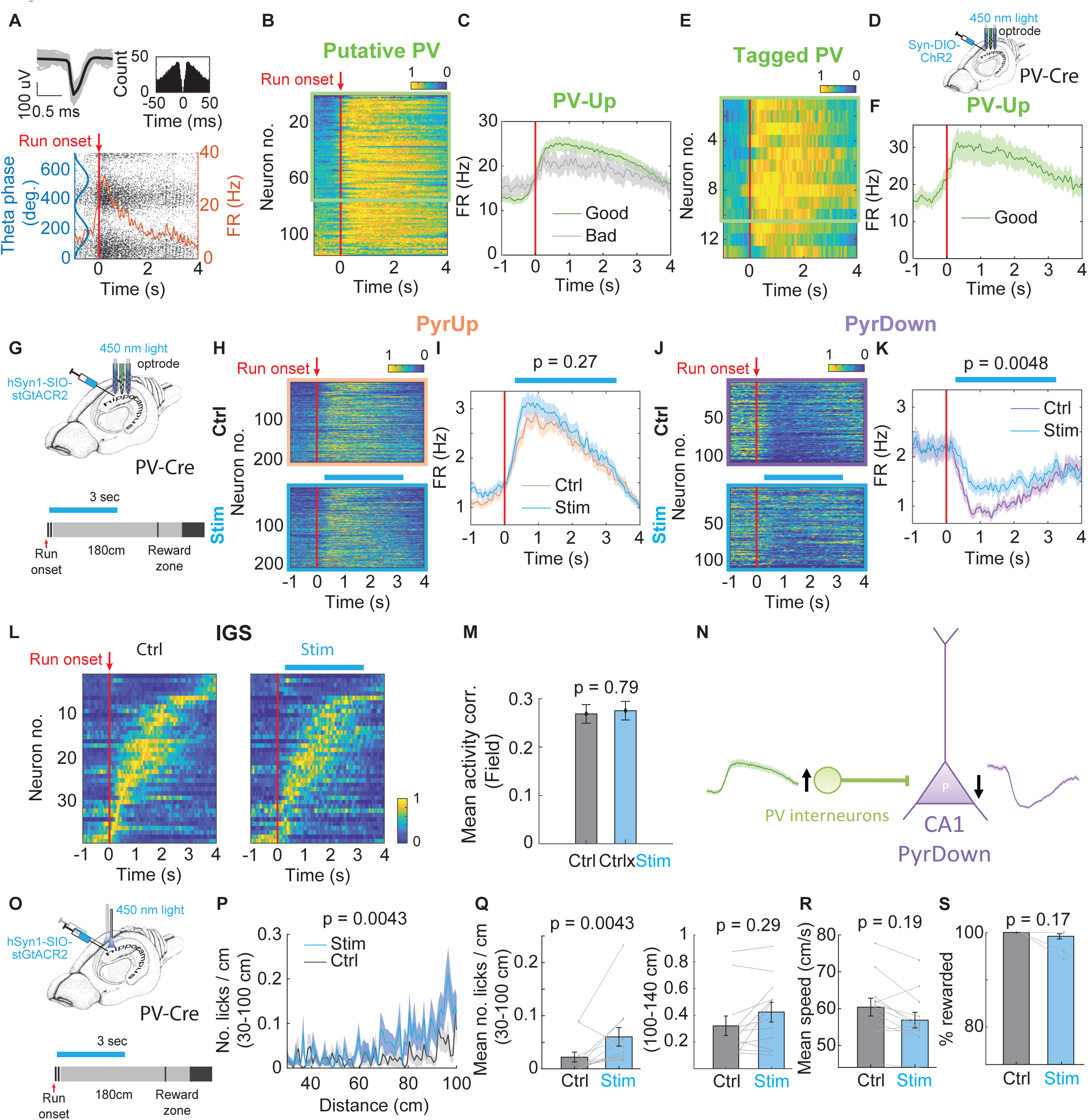
CA1 parvalbumin-positive interneurons regulate PyrDown response and the start of integration. (A) Spike waveform (top-left) and auto-correlogram (top-right) of a putative PV interneuron. Raster plot (bottom) showing spike theta phases (black dots) and firing rate as a function of time (orange) for the same cell. (B) Normalized firing rate heatmaps of all the interneurons belonging to the putative PV interneuron cluster, ordered by their firing rate ratio FR_aft_/FR_bef_ around run onset. The green box denotes PV-Up neurons whose FR_aft_/FR_bef_ > 3/2. (C) Firing rate profile averaged across all PV-Up neurons broken down into good and bad trials. Relative FR change: good trials: 0.33±0.012, bad trials: 0.16±0.021, p=1.02e-8. (D) Schematic for the opto-tagging experiment. (E) Same as in (B), for interneurons identified as PV-positive from the tagging experiments and at the same time belonging to the putative PV interneuron cluster (10/13 neurons with FR_aft_/FR_bef_ > 3/2). (F) Bottom, same as in (C), but for tagged PV-Up neurons highlighted within the green box in (E). Bad trials are not shown because the number of bad trials is less than 15 trials in most of these recordings. (G) Experimental setup for inactivating PV interneurons. Light stimulation is applied with an optrode for 3 seconds starting after run onset in a subset of trials. (H) Normalized firing rate heatmaps of all PyrUp neurons during control (top) and stimulation trials (down). For details, see Figure 4H. (I) Firing rate profile averaged across PyrUp neurons. Mean FR between 0.5-2s: control: 2.64±0.16Hz, stimulation: 2.90±0.17Hz, p=0.27 (two-sample t-test). (J-K) Same as in (H-I), for PyrDown neurons. For details, see Figures 4H and 4I. Mean FR between 0.5-2s: 0.97±0.091Hz, stimulation: 1.43±0.13Hz, p=0.0048 (two-sample t-test). (L-M) Same as in Figures 4L and 4M, for inactivating PV interneurons. Mean activity corr.: control: 0.27±0.019, stimulation: 0.28±0.019, p = 0.79. (N) A circuit diagram potentially illustrates how PV inactivation impacts PyrDown response. (O-S) Same as in Figures 4O-4S, for inactivating PV interneurons, including 4 animals, 12 recordings. (Q) left: control: 0.022±0.0093, stimulation: 0.060±0.073. right: control: 0.32±0.073, stimulation: 0.42±0.074. (R) control: 60.40±2.42cm/s, stimulation: 56.87±2.14cm/s. (S) control: 100.00±0.00%, stimulation: 99.18±0.56%.

In this cluster, a majority of neurons (75/114 neurons, Figures 6B and 6C), including most of the tagged cells (10/13 neurons, Figures 6E and 6F), exhibited a rapid increase in activity around run onset (PV-Up neurons). In bad trials, the PV-Up response was weakened without a change in mean firing rate (Figure 6C, mean FR: good trials: 20.16±1.29Hz, bad trials: 18.27±2.15Hz, p=0.47).

We further investigated the timing of the PV-Up response relative to PyrUp and PyrDown responses. Both PV-Up and PyrUp neurons showed concurrent increases in normalized firing rates around run onset, with PV-Up neurons leading PyrUp neurons in time (Figure S4H left, time to reach 90% of maximum firing rate within [-1 1.5]s window (t_90%_): PyrUp: 0.77±0.01s, PV-Up: 0.50±0.04s, p=7.53e-09). In contrast, PyrDown response started to decrease only after the initial rise of PV-Up response and reached its minimum after the peak of PV-Up response (Figure S4H right, t_90%_: PyrDown: 0.60±0.01s, p=6.62e-04 compared with PV-Up). These results imply that PV-Up neurons may preferentially inhibit PyrDown neurons.

### Parvalbumin-positive interneurons are necessary to start distance counting at run onset

To determine if PV interneurons predominantly evoke inhibition on PyrDown neurons, we used stGtACR2 to silence CA1 PV interneurons immediately after run onset for 3 seconds (Figure 6G, Methods)^31^. This manipulation effectively silenced some putative PV cells (Figures S4K and S4L).

Silencing PV interneurons caused a significant elevation in the PyrDown response after run onset (Figures 6J and 6K, 3 animals, 6 recordings, 205/528 neurons), comparable with the results in bad trials (Figure 3I). However, neither the PyrUp response (Figures 6H and 6I, 108/528 neurons) nor IGSs (Figures 6L and 6M, 39/528 neurons, mean FR: control: 1.96±0.18Hz, stimulation: 2.23±0.23Hz, p=0.33) showed significant changes.

To assess behavioral consequences, we bilaterally silenced PV interneurons after run onset in a separate cohort of animals (Figure 6O). In stimulation sessions, there was a significant increase in early licks within 100 cm (Figures 6P and 6Q left), but no changes beyond 100 cm (Figure 6Q right). That is, the impairment was mostly confined to shortly after run onset. The percentage of rewarded trials (Figure 6S) and mean running speed (Figure 6R) did not change. Control experiments ruled out that the effects were due to laser light alone (Figures S5A-S5E, S5K-S5O; Methods).

The timing of the behavioral impairment occurring shortly after run onset implies that the animals cannot accurately start integration. Thus, we posit that PV interneurons are necessary for shutting down PyrDown neurons at run onset (Figure 6N) and initiating integration over distance or time at the beginning of each trial.

## DISCUSSION

In this study, we uncovered two functional subpopulations of CA1 pyramidal neurons, PyrUp and PyrDown neurons, that offer a novel mechanism for encoding elapsed time or traveled distance. PyrUp neurons exhibited a collective increase in activity to mark the start of integration, followed by individual neurons ramping down at varying rates that encode the passage of time. Conversely, PyrDown neurons initially decrease their activity before gradually ramping up toward the reward. Both subpopulations potentially use their response at the integration start as a reference point for the subsequent distance or time encoding.

We also identified two parallel CA1 circuits involving local inhibitory interneurons that regulate individual subpopulations and influence integration accuracy. Somatostatin-positive interneurons preferentially modulated the PyrUp response and IGSs and were necessary for accurate time or distance integration. In contrast, parvalbumin-positive interneurons primarily regulated the activity of PyrDown neurons and were necessary for correctly initiating integration.

These findings support the hypothesis that SST interneurons define an integration window, facilitating PyrUp response and IGSs during integration; while PV interneurons generate a reset, shutting down PyrDown neurons and reinitiating integration. The coordinated interaction between these interneuron types and their corresponding pyramidal subpopulations orchestrates different aspects of the time or distance integration.

### PyrUp and PyrDown responses provide a novel coding mechanism for time or distance

PyrUp and PyrDown neurons provide a novel two-phase mechanism for encoding time or distance. In PyrUp neurons, the first phase marks the initiation of integration, characterized by a synchronized rise in firing until peaking at similar times across the population shortly after run onset (Figure 3K). The second phase encodes time or distance, as the individual neurons’ firing rates diverge due to the heterogeneous decay rates after the peak. Larger divergences in the firing rates correspond to longer elapsed times.

Based on these observations, we propose that the PyrUp neurons first generate a reference signal through the synchronized rise to peak, which then serves as the starting point for subsequent encoding of time or distance via divergent decay. PyrDown neurons follow a similar two-phase coding mechanism (Figures 3M–3N). This coding strategy is distinct from the sequential firing patterns formed by neurons with place fields and IGFs. It highlights a novel mechanism for temporal and spatial encoding, a critical component during the moment-to-moment experience encoding.

### A small proportion of neurons exhibit IGFs in the PI task

Firing fields in hippocampal neurons have been well-established in the literature. Prior studies suggest that in environments with minimal sensory cues or virtual reality settings, hippocampal neurons show reduced spatial tuning. Correspondingly, a relatively small proportion of neurons express firing fields^35–37^. However, in cue-poor or cue-absent environments, the generation of firing fields (IGFs) is influenced by cognitive and memory demands. The prevalence of IGSs typically increases under conditions requiring higher demands^6,8–10,20,38–40^.

Consistent with these prior findings, in our PI and immobile tasks, which are less demanding but still require animals to track distance or time, only a small percentage of neurons expressed IGFs (Figure 2G). In contrast, IGFs were absent in the passive task that does not require integration, echoing previous observations^6,20^. Moreover, our results align with earlier evidence^10,41^ that IGSs became impaired when animals failed to estimate distance or time accurately. Together, these data support the idea that tasks requiring higher cognitive demand or encoding accuracy tend to recruit a larger proportion of neurons to express IGFs.

### PyrUp/PyrDown neurons are not simply “wide” firing fields

In this study, we define PyrUp and PyrDown neurons as pyramidal neurons that exhibit a clear change in firing rate around run onset — a criterion that does not exclude neurons with IGFs. Some pyramidal neurons classified as PyrUp or PyrDown neurons exhibited IGFs around the run onset or near the reward zone. However, in the PI task, only ∼10% of pyramidal neurons were identified as having firing fields, whereas PyrUp and PyrDown neurons collectively accounted for ∼56% of all recorded pyramidal neurons. Excluding the subset of PyrUp and PyrDown neurons with IGFs from our analysis did not alter our core findings.

We would like to note that PyrUp and PyrDown neurons without IGFs are unlikely to be simply neurons with wide firing fields. Neurons with IGFs exhibited significantly stronger phase precession, a key characteristic of firing fields^21,22^, than those without (Figure S1G). Moreover, classifying all PyrUp and PyrDown neurons as firing-field cells—regardless of their tuning specificity—would represent ∼56% of the entire pyramidal population. This proportion is notably higher than usually reported for neurons with firing fields in similar cue-poor contexts^35,36^. In summary, while PyrUp/PyrDown neurons and neurons with IGFs are not mutually exclusive, most PyrUp and PyrDown neurons do not meet commonly accepted criteria of firing field in hippocampal research, including tuning specificity and robust phase precession.

Even if all PyrUp and PyrDown neurons were classified as having IGFs, our central conclusion remains valid: PyrUp/PyrDown neurons can encode time and distance through a mechanism distinct from the sequential patterns formed by neurons with firing fields. While neurons with firing fields encode time or distance through specific tuning around particular time points or distances, PyrUp and PyrDown neurons use a population-level two-phase coding mechanism that does not rely on tuning specificity. This novel coding mechanism provides insights into alternative coding mechanisms within the hippocampal network.

### The relationship between PyrUp and PyrDown responses and running speed

Although PyrUp and PyrDown responses in the PI task correlate with locomotion, they cannot be fully explained by running alone. Several lines of evidence support this conclusion. First, PyrUp and PyrDown neurons exhibited context-dependent responses, showing significantly stronger responses at the onset of trial-start runs than spontaneous runs occurring during a trial. This result remained consistent even after matching running speed (Supplementary Figures 3A-3C). Second, PyrUp and PyrDown populations were also observed in an immobile task—where animals stayed still while estimating a 4-second time interval—demonstrating that these responses can arise without locomotion (Figures S3D–S3J). Third, PyrUp neurons began ramping up their activity before run onset (Figures 3G and 3K), indicating that their activity was not purely a reflection of running speed. In summary, although running speed may influence PyrUp and PyrDown responses, these responses extend beyond simple movement-related signals.

### Differential role of CA1 SST and PV interneurons in internally generated neuronal dynamics

Our results underscore the distinct roles of CA1 SST and PV interneurons in shaping internally generated dynamic patterns produced in the absence of changing sensory cues, including PyrUp and PyrDown responses and IGSs.

First, our data support that SST interneurons define a window of inhibition that begins with a gradual increase in firing around run onset and tapers off as the reward zone gets close. Within this window, these interneurons exert stronger control over PyrUp neurons and IGSs (Figure 4). Counter-intuitively, silencing SST interneurons after the run onset inhibited the PyrUp response, implying that SST interneurons disinhibited PyrUp neurons by suppressing other inhibitory interneurons and favoring CA3 inputs. Behaviorally, inactivating SST interneurons impaired distance or time estimation when performed after run onset but not before the reward zone. These results support the role of SST interneurons in sustaining continuous integration.

Our data also suggest that SST interneurons primarily affect internally generated neuronal dynamics, instead of sensory-driven responses. When visual cues were introduced into the cue-constant segment in the PI task (cue-rich task), inactivating SST interneurons after run onset no longer yielded significant changes in neuronal sequences or PyrUp response.

Second, PV interneurons may generate a reset signal by rapidly ramping up their firing around run onset, which leads to the synchronized shutdown of PyrDown neurons (Figure 6). Behaviorally, inactivating PV interneurons after run onset affected integration accuracy only at the start. These results align with the hypothesis that PV interneurons reset the distance or time counter, starting a new period of integration.

While PV interneurons may ensure a correct start of integration, they are unlikely to be the only circuit elements involved. The start of integration is likely encoded in diverse populations of neurons across regions providing inputs to CA1, including other hippocampal areas and the entorhinal cortex. This idea is supported by the mild behavioral deficit observed during PV interneuron inactivation after run onset.

### Overlap of circuits that initiate integration and delineate ongoing experience into discrete units

Our data suggest that in the PI task, the run onset at the trial’s start may serve as the start of neural computation for distance or time integration. First, IGSs started at run onset in the PI task. This result agrees with a previous report that IGSs can begin following the onset of locomotion^38^. Second, PyrUp and PyrDown neurons exhibited strong firing rate changes around run onset at the trial’s start, but not around spontaneous running onset. However, run onset is not the only cue that can signal the start of integration. For instance, in our immobile task, the last lick appears to serve a similar role as an initiating cue.

More generally, animals can use salient sensory^11,42^ or motor cues^38^ in the trial structure to initiate integration and delineate continuous experience into individual units. These results resonate with studies on human memory encoding, which suggest that the continuous stream of experience can be segmented into discrete units by detecting event boundaries^43,44^. The salient sensory or motor cues described in past and current studies may act as such boundaries, with the process of integration within each trial comparable with integrating ongoing experience within a discrete unit.

Overall, the CA1 neural circuit motifs that we identified may contribute to the computation performed in the process of detecting event boundaries and encoding continuous experiences into individual units.

## ACKNOWLEDGMENTS

We thank T. Harris, E. Moser for comments on the manuscript; L. Abbott, E. Schuman and D. Fitzpatrick for discussions; S. Sawtelle, B. Wisnicki and X. Zhao for help with the virtual reality setups. M. Klement, N. Daniel and machine shop at MPFI for making mechanical parts for the experimental setups; J. Wells and ARC at MPFI for taking care of animals. This work was funded by Max Planck Society and Max Planck Foundation, NIH R01 NS119503.

## AUTHOR CONTRIBUTIONS

Y.W. conceived the project. Y.W. and R.H. designed experiments (with input from D.P.). R.H. performed electrophysiological experiments. D.P., Y.W, and X.Z. performed behavioral experiments with optogenetics. Y.W. and R.H. analyzed data. Y.W., B.M., and R.H. discussed the results and wrote the manuscript, with contribution from all the authors.

---

**Figure S1.**
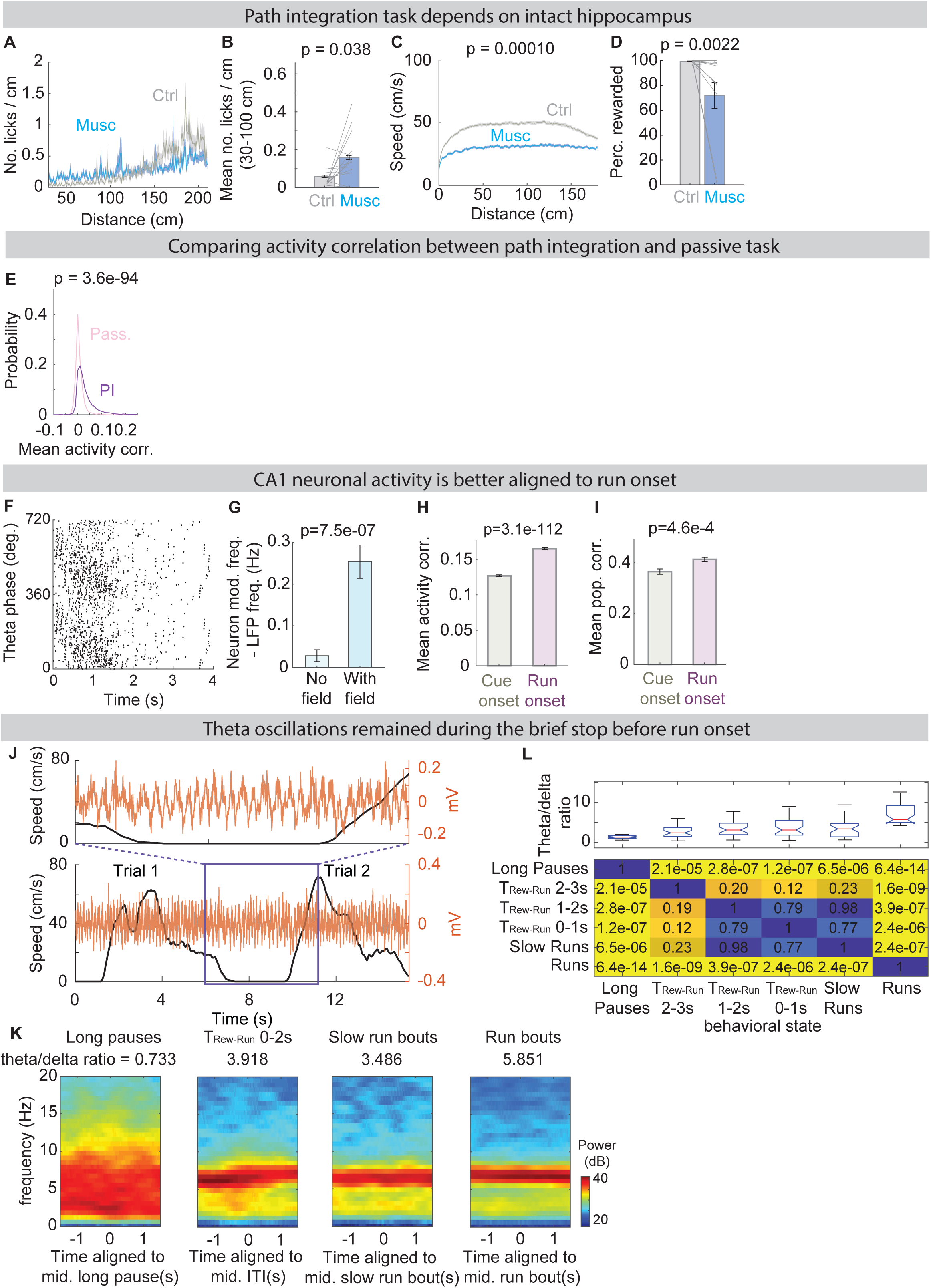
Internally generated sequence (IGS) starts at run onset in the PI task, related to Figures 1 and 2. (A-D) The hippocampus dependence of the PI task is assessed by muscimol infusion experiments targeting CA1. (A) Mean lick histogram binned by distance during the control (grey) and muscimol (blue) sessions (9 animals, 10 recordings). (B) Mean number of licks/cm in the early portion of the trial. (C) Average speed curves over distance. (D) The percentage of trials where the animal received a reward. (E) Distribution of mean trial-by-trial activity correlations of all the pyramidal neurons recorded in the passive (536 neurons) and PI tasks (7828 neurons). (F-G) Phase precession in neurons with IGFs. (F) A raster plot showing the theta phase change over time for the example pyramidal neuron in Figure 1F right that has an IGF and demonstrates phase precession. The spikes are aligned with run onset. (G) Pyramidal neurons with fields display significant phase precession relative to those without fields after aligning with run onset (7178 neurons). Phase precession is calculated as the difference between the oscillating frequency of individual neurons and the local field potential theta oscillation frequency. (H-I) Alignment of all pyramidal neurons. (H) Mean activity correlation aligned with cue vs. run onset for all pyramidal neurons (7828 neurons). (I) Mean population correlation aligned with cue vs. run onset for all pyramidal neurons per recording (102 recordings). (J-L) Theta oscillations are unchanged when an animal initiates running at the beginning of each trial after stopping to consume reward at the end of the last trial. (J) An example showing that theta oscillations persist during the brief stop before run onset. Black: running speed. Red: raw local field potential trace. Top: zooming in the segment inside the purple box. (K) Averaged multitaper spectrograms and theta/delta ratios aligned with the midpoints of long pauses (left), trials where the interval between the last trial’s reward delivery and the current run onset (T_Rew-Run_) is between 0-2s (middle left), slow running (speed 5-20 cm/s, middle right) or running bouts (right) in one example recording. (L) Top: distribution of theta/delta ratios for different behavioral states pooled across recordings (13 animals, 25 recordings). Bottom: p-value matrix comparing theta/delta ratios across behavioral states using pairwise Kruskal-Wallis tests. There is no significant change during the pause between trials and during slow running (speed 5-20 cm/s).

**Figure S2.**
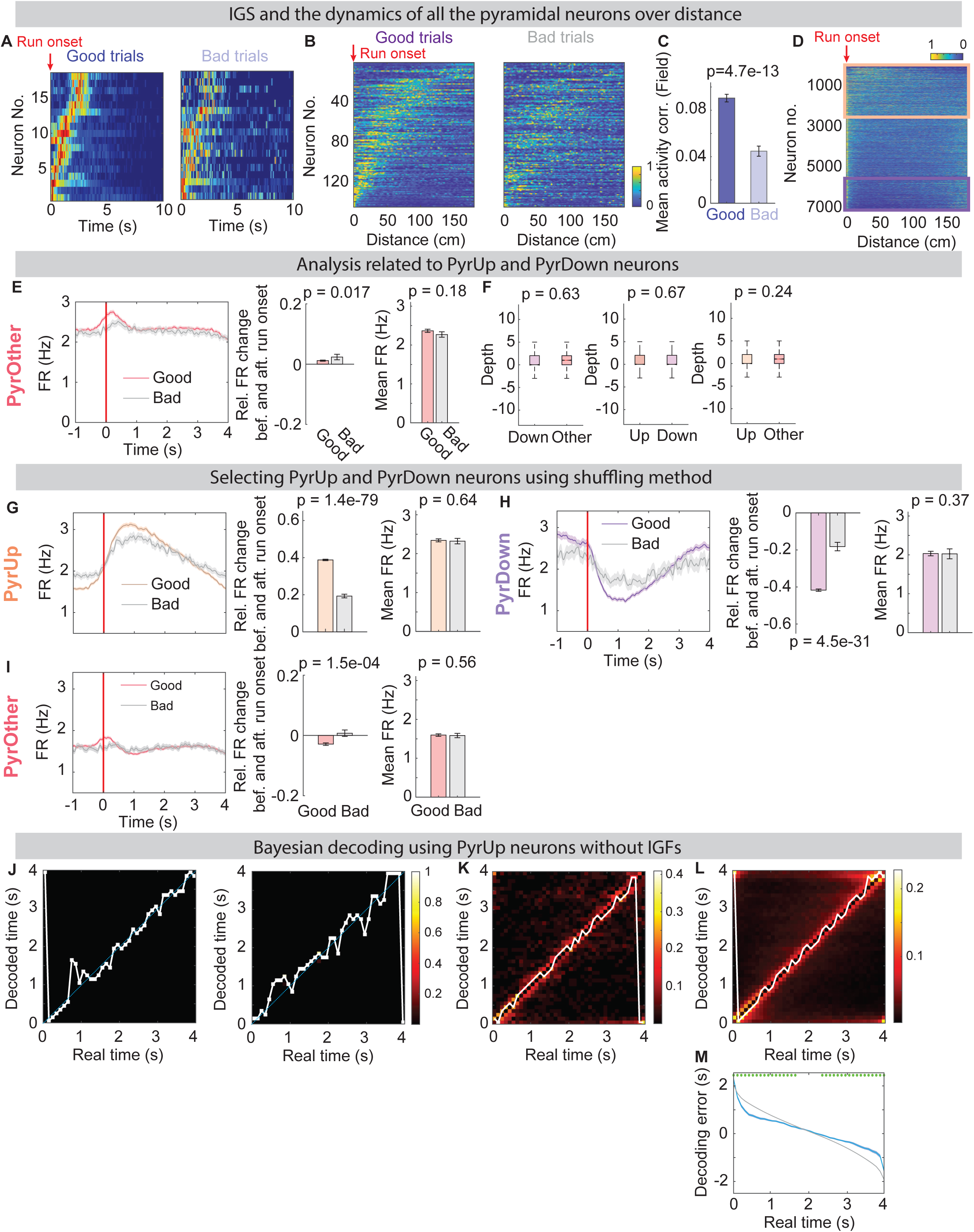
Pyramidal neurons display distinct responses around run onset, related to Figure 3. (A) An example IGS from the same session as in Figure 1G averaged across good (left) and bad (right) trials. (B) The same sequence as in Figure 3C, plotted over distance. (C) Same as in Figure 3D, but the correlation is calculated based on the firing rate profiles over distance. (D) Same as in Figure 3E, plotted over distance. (E) Firing rate profile averaged across all PyrOther neurons (2998 neurons), broken down into good and bad trials (Left). Relative firing rate change around run onset (Middle), and overall mean firing rate calculated between −1s and 4s (Right). (F) Depth distribution comparing all PyrDown and PyrOther neurons (left), PyrUp and PyrDown neurons (middle), and PyrUp and PyrOther neurons (right), respectively. (G) Normalized firing rate profile for PyrUp neurons identified by shuffling method (2943 neurons, see Methods) in good and bad trials (left). Relative firing rate change around run onset (middle) and overall mean firing rate (right). (H) Same as in (G), for PyrDown neurons identified by shuffling method (923 neurons). (I) Same as in (G), for PyrOther neurons identified by shuffling method (3154 neurons). (J-M) Bayesian decoding using PyrUp neurons without IGFs. (J) Decoding results of two single trials. Color bar indicates posterior probabilities. White line denotes the decoder’s most confident estimation. (K) Decoding results of a single recording session, the same session as in (J). (L) Decoding results averaged across all recording sessions (23 animals, 70 recordings). (M) Decoding error determined using tenfold cross-validation (blue) compared with the shuffled data (grey), averaged across all recording sessions. Green dots highlight the time points where the decoding error is significantly different from shuffled data.

**Figure S3.**
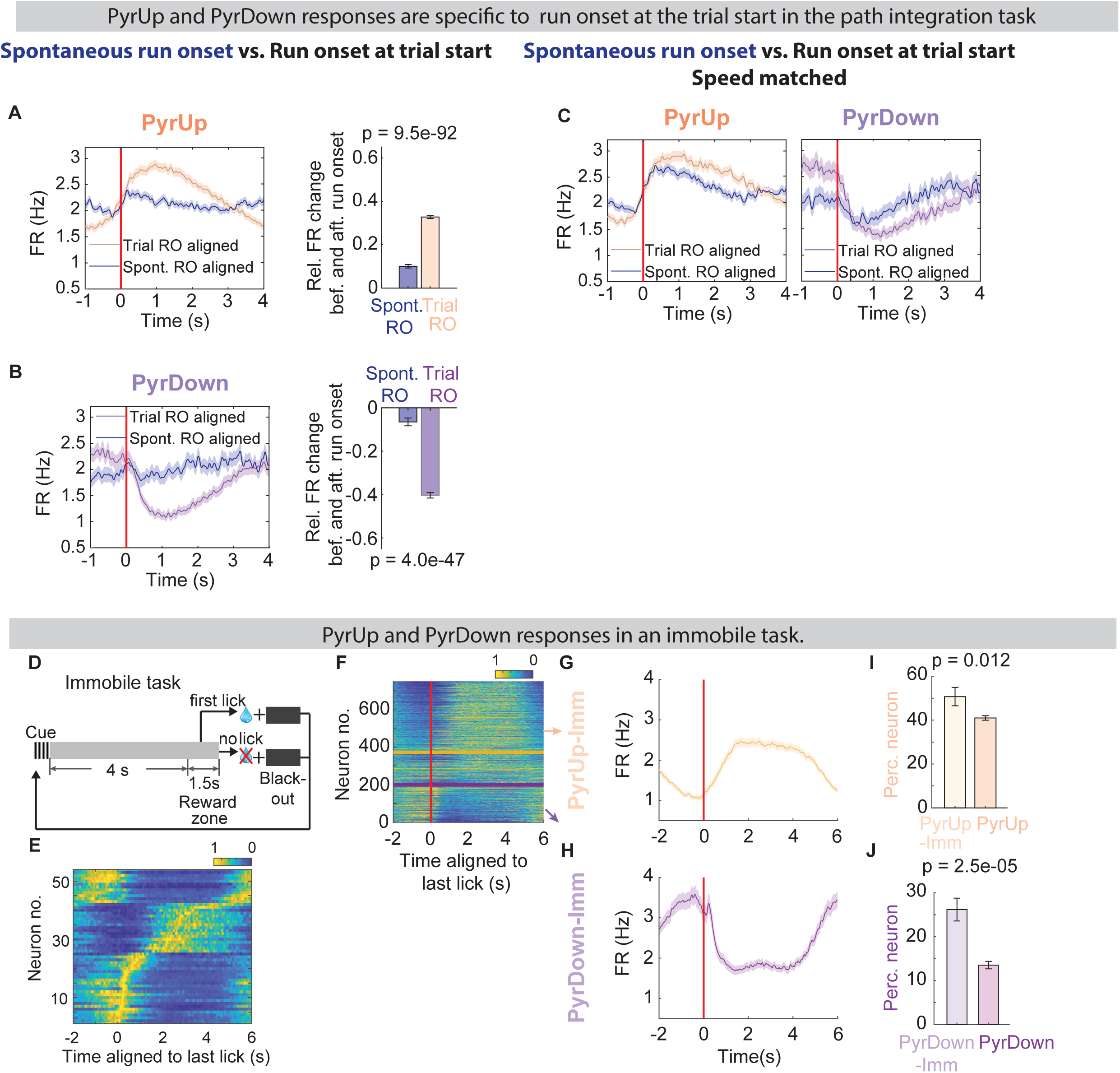
Pyramidal neurons display distinct responses around the start of integration, related to Figure 3. (A-C) PyrUp and PyrDown responses are specific to run onset at the trial start in the PI task. (A) Left: After realigning with spontaneous run onset (SRO), the firing rate profile averaged across PyrUp neurons (dark blue) shows smaller changes than after aligning with the trial-start run onset (TRO). Recordings with at least 15 spontaneous run bouts were included in the analysis. Right: Relative firing rate change around run onset. At SRO: 0.10±0.008, at TRO: 0.33±0.0069. (B) Same as in (A), for PyrDown neurons. At SRO: −0.065±0.018, at TRO: −0.40±0.012. (C) Similar to (A) and (B), after selecting recordings with matching running speed between spontaneous runs and runs from the trial start (see Methods). (D-J) PyrUp and PyrDown responses in an immobile task. (D) Task schematic for the immobile task. (E) IGS is present after aligning the neuronal activity with the last lick of the previous trial (53/748 neurons). (F) Normalized firing rate heatmaps of all pyramidal neurons ordered by their firing rate ratio FR_aft_/FR_bef_ (FR_bef_: the mean firing rate (FR) before last lick (−1.5->-0.5s), FR_aft_: FR after last lick (0.5->1.5s), 0s corresponds to the last lick). PyrUp (377 neurons) and PyrDown (199 neurons) neurons identified by shuffling method are separated by orange and purple lines, respectively. (G) Normalized firing rate profile averaged across all PyrUp neurons. (H) Same as in (G), for PyrDown neurons. (I) The percentage of PyrUp neurons identified in the immobile task (yellow) is higher than that identified in the PI task (orange). (J) Same as in (I), for PyrDown neurons.

**Figure S4.**
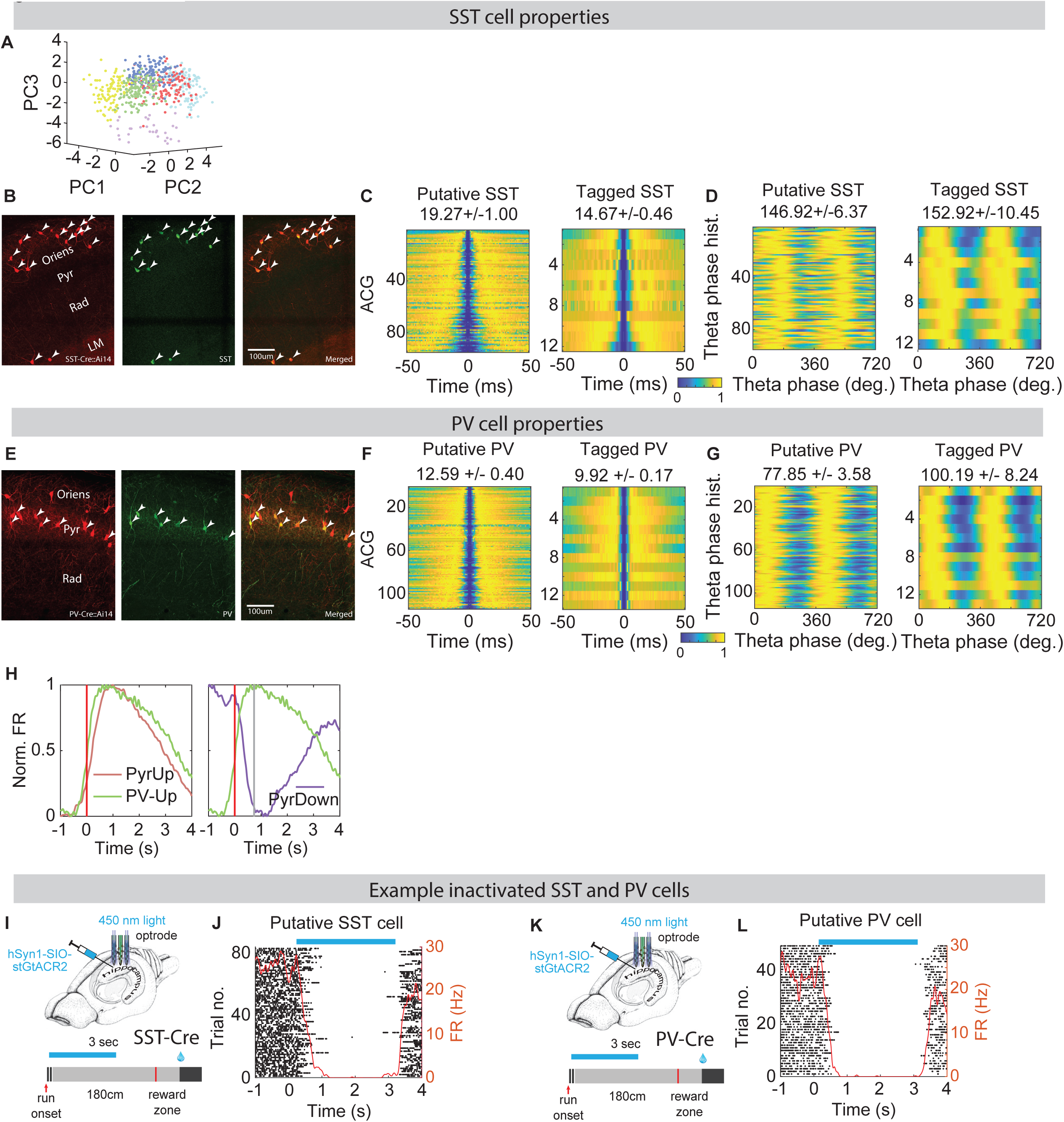
Properties of SST and PV interneurons, related to Figures 4 and 6. (A-D) SST interneurons identification through optogenetic tagging. (A) A three-dimensional view of the 6 interneuron clusters in the PCA space. (B) Confocal images of CA1 from an SST-CrexAi14 mouse. From left to right, tdTomato expression, SST immunostaining (GFP), and overlaid images. Arrows point to cells that are co-labeled for tdTomato and GFP. (C) Auto-correlograms (ACGs) of putative (left, as in Figure 4B) and tagged SST cells (right, as in Figure 4E). On the top shows the mean+/− sem of the ACG peak time averaged across all cells. (D) Theta phase histograms of putative (left) and tagged SST cells (right). On the top shows the mean+/− sem of the peak theta phase averaged across all cells. (E-G) PV interneuron identification through optogenetic tagging. (E) Confocal images of CA1 from a PV-CrexAi14 mouse. From left to right, tdTomato expression, PV immunostaining (GFP), and overlaid images. (F) Auto-correlograms (ACGs) of putative (left, as in Figure 6B) and tagged SST cells (right, as in Figure 6E). (G) Theta phase histograms of putative (left) and tagged SST cells (right). (H) Averaged normalized firing rate profiles of PyrUp and PV-Up neurons (left), or PyrDown and PV-Up neurons (right). The peak of PV-Up neurons’ activity is denoted by the grey bar (right). (I-J) Optrode-mediated inactivation of SST interneuron activity at run onset. (I) From Figure 4G, experimental setup. (J) An example putative SST cell that is effectively inactivated by light. (K-L) The same as (I-J), for PV interneurons. (K) From Figure 6G.

**Figure S5.**
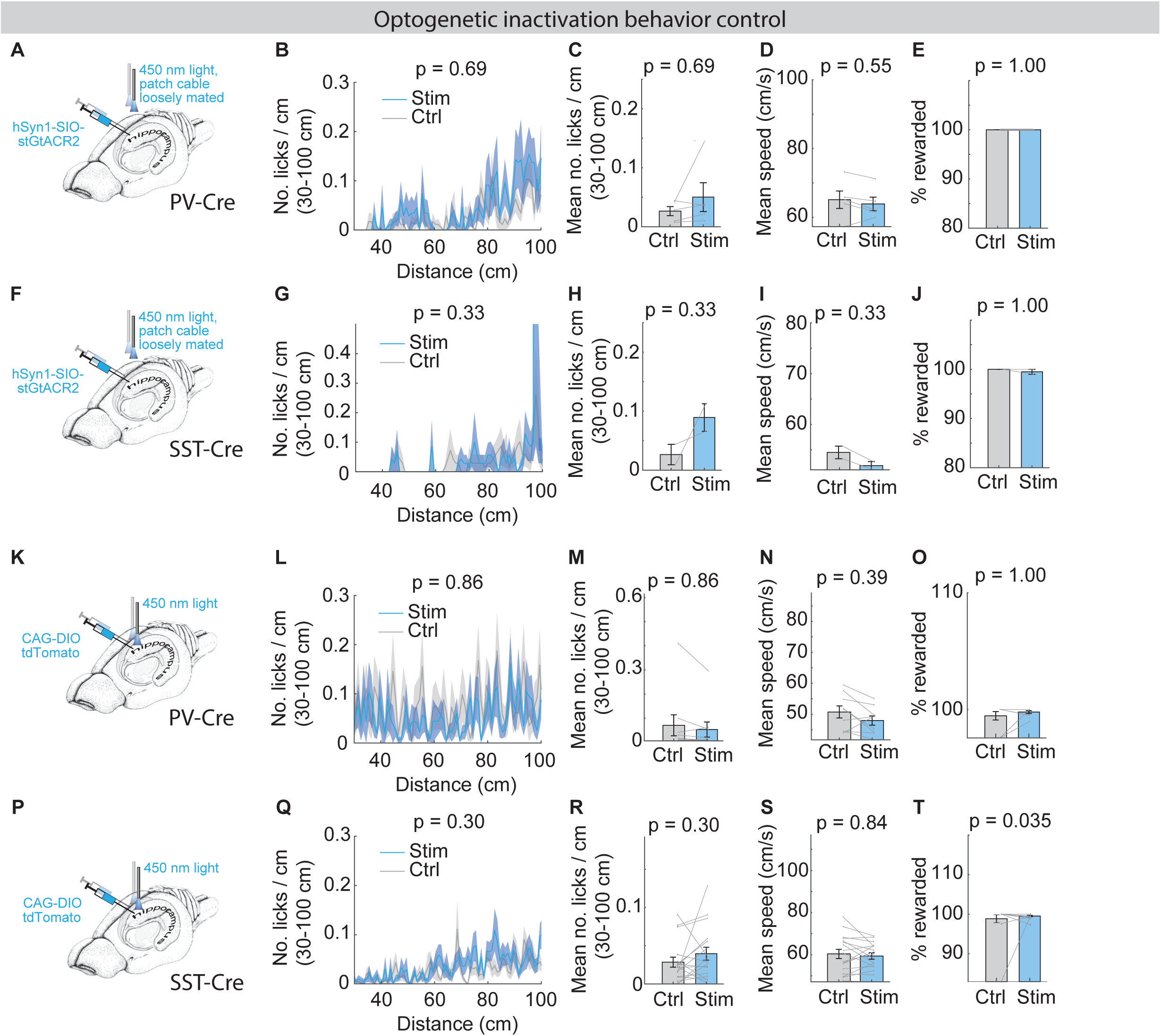
Control experiments for bilateral optogenetic inactivation, related to Figures 4 and 6. (A-E) Control experiments for optogenetic inactivation of PV interneurons (3 animals). (A) Experimental design. Light delivery protocols are identical to previous experiments, but the patch cables were adjusted not to be in contact with the implanted fiber optic cannulas, precluding light delivery into the brain. (B) Lick histogram for the first 100 cm for control and stimulation sessions. Reported p-value is for the mean number of licks in this same period. (C) Mean number of licks/cm calculated based on (B). (D) Mean running speed. (E) Percentage of rewarded trials. (F-J) The same as (A-E), for control experiments for inactivation of SST interneurons (2 animals). (K-O) The same as (A-E), for control experiments using interneurons infected with tdTomato (2 animals). (P-T) The same as (F-J), for control experiments using interneurons infected with tdTomato (4 animals).

## METHODS

### EXPERIMENTAL MODEL AND SUBJECT DETAILS

#### Mice

This study was based on both male and female mice (age > 8 weeks). Male mice were preferentially used in the running tasks because they were found to exhibit more consistent running behavior. We used four mouse lines: C57Bl/6J (JAX #000664), PV-IRES-Cre (JAX #017320) (Hippenmeyer et al., 2005), SST-IRES-Cre (JAX #013044) (Taniguchi et al., 2011), and Ai14 (JAX # 007914) (Madisen et al., 2012). All procedures were in accordance with protocols approved by the Institutional Animal Care and Use Committee at Max Planck Florida Institute for Neuroscience. Mice were housed in a 12:12 reverse light: dark cycle and behaviorally tested during the dark phase.

### METHOD DETAILS

#### Virtual reality setups

For the virtual reality (VR) setups, a small treadmill (Janelia Design) was positioned in front of two visual displays, such that after head-fixation, the eyes aligned with the two visual displays. Customized software was written using Psychtoolbox to display the virtual environment. An Arduino control board was used to control when to display the visual stimuli. For the small treadmill, the animal’s speed was measured using an encoder attached to the back wheel axis. A microprocessor-based (Arduino) behavioral control system (the miniBCS board, designed at Janelia) interfaced with a MATLAB graphical user interface controlled the trial structure, the water valve, and the encoder. A separate lick port detector (designed at Janelia) was used to convert a touch on a metal lick port into a digital pulse and to send the information to the miniBCS board. In another version of the VR setups, instead of using two monitors, a curved screen was used where the virtual reality environment was projected onto the curved screen using a projector (AKASO Mini Projector, a design kindly shared by Christopher Harvey’s lab at Harvard University). In this case, customized software written with Unity was used to display the virtual environment.

In addition, a separate microprocessor (Arduino) interfaced with a customized MATLAB graphical user interface was used to operate the laser or laser diode on-off for the optogenetic perturbation experiments and to control the closed-loop manipulation of PVand SST interneurons. Behavioral data were monitored and recorded as well using MATLAB functions.

#### Behavioral training

Before any surgery was performed, running wheels were added to the home cages. At 3–5 days after the headbar or fiber implantation, mice were placed on water restriction (typically 0.8-1 ml per day). After each training or recording session, mice were supplemented with additional water to ensure the amount of water intake per day. After habituating the animals to the treadmill for at least 1-2 days, animals were trained to run head-fixed on the treadmill.

In the passive task, each trial started with a grey-colored visual cue. After the animal ran for 180 cm while this visual stimulus stayed constant, a water reward was automatically delivered. After that, an inter-trial-interval (ITI) started while the visual stimulus kept constant as before. The next trial started at least after 0.5s and until the animal’s speed has exceeded 30 cm/s for more than 0.3s and the time since the last lick has exceeded 0.3s. Therefore, there was a random ITI of at least 0.5s until the next trial started. It took about 1-2 weeks for the animals to run smoothly on a treadmill and to perform this task.

In the PI task, a trial started with a visual grating stimulus that lasted either 0.5s or 1s, and stayed the same within each session. After that, a grey-colored stimulus turned on. The animal was required to run for a fixed distance (180 cm) while this visual stimulus stayed constant, and then it must lick within a 40 cm (in some sessions, it is 80 cm) unmarked reward zone to receive a water reward. The first lick in the reward zone triggered a reward, and simultaneously the screens turned black for either 0.5s or 1s (kept constant within each session). If no lick occurred in the reward zone, the screens turned black to signify the error for 0.5-2s (kept constant within each session). It took about 2 weeks for the animals to learn this task.

In the immobile task, the animal sat within a plastic tube^29^. Each trial started with a 1s visual grating stimulus, which was then followed by a grey-colored stimulus that lasted 4s. After that, there was a reward zone lasting 2s. If the animal licked within the reward zone, a drop of water reward was triggered, and the screens turned black for a random time between 0.5-2.5s. If the animal failed to lick within the reward zone, the screens turned black for a random time between 0.5-2.5s. To train the animal to perform this task, the animal was first habituated to stay in the tube, and then head fixation was gradually applied. This task took about 2 weeks to train.

#### Virus injection, headbar, cannula and fiber implantation

Adult mice (2-4 months of age) underwent aseptic stereotaxic surgery to implant a custom lightweight 3D printed headbar under isoflurane anesthesia (2-3% for induction, 1%–1.5% for maintenance). Buprenorphine SR LAB (0.5 mg/kg, SC) or buprenorphine (0.1 mg/kg, SC), and Meloxicam SR (5 mg/kg, SC) was administered immediately after the surgery. For acute extracellular recordings, after training to perform behavior tasks, craniotomies were performed which were centered around antero-posterior 2.1 mm from bregma, mediolateral +/−1.7 mm from the midline to target dorsal CA1 region. Meanwhile, a ground wire was attached to the headbar, and a thin layer of silver paint (GC ELECTRONICS 22-0023-0000) was applied to the surface of the headbar for noise reduction during the recording. The skull was covered using KwikSil (World Precision Instruments) and was only removed during recordings.

For extracellular recordings with optogenetic manipulation of interneurons, AAV1_hSyn1-SIO-stGtACR2-FusionRed (Addgene 105677, 2.1e+12 after 10 fold dilution) was used for optogenetic inactivation, and AAV5-Ef1a-double-floxed-hChR2(H134R)-mCherry-WPRE-hGHpA (Addgene 20297, titer 1.4e+13) was used for optogenetic activation. 50 nl virus per hemisphere was injected bilaterally in the CA1 of PV-IRES-Cre and SST-IRES-Cre mice. The injection was done at least 3 days before the headbar implantation, and at least 3 weeks before the recording. The following coordinates were used for viral injections: 2.1 mm from bregma, mediolateral +/−1.7 mm from the midline and 1.24 mm dorsoventral from the brain surface. The injection system comprised a pulled glass pipette (broken and beveled to 15-20 μm inside diameter; Drummond, 3-000-203-G/X), backfilled with mineral oil (Sigma). A fitted plunger was inserted into the pipette and advanced to displace the contents using a manipulator (Drummond, Nanoject II or Nanoject III). Retraction of the plunger was used to load the pipette with the virus. The injection pipette was positioned onto a Kopf manipulator.

For optogenetic experiments without extracellular recordings, the viral injection procedure was the same as described in the last paragraph. At least 3 days after the viral injection, optical fibers (core diameter of 200 um) were chronically implanted bilaterally to target the CA1 region using the following coordinates: 2.1 mm from bregma, mediolateral +/−1.7 mm from the midline and 0.9 mm dorsoventral from the brain surface.

For muscimol infusion experiments, guide cannulas (26 Gauge) were implanted bilaterally above the CA1 region (coordinates: 2.1 mm from bregma, mediolateral ±1.7 mm from the midline and 1.1 mm dorsoventral from the brain surface) during the head bar surgery. Dummy cannulas (33 Gauge) of the same length as the guide cannulas were inserted into the guide cannulas.

#### Optogenetics without extracellular recordings

For each optogenetic session, a control session (∼40 trials) was performed. It was then followed by a stimulation session (∼100 trials in total), where the photostimulation was deployed every third trial. After that, there was a post stimulation control session (∼40 trials). To prevent mice from distinguishing photostimulation trials from control trials through stimulation light, a masking blue light (470 nm LEDs (Thorlab)) was on throughout the sessions. For both stGtACR2 and ChR2, we used a 473 nm laser (Ningbo Lasever Inc.). The laser power used was <= 5 mW. During the stimulation, 100 Hz light pulses were used to reduce the heating effect of a constant laser light. The duty cycle was varied to adjust the light power.

For interneuron inactivation using stGtACR2, the stimulation lasted 3 seconds. The stimulation can be triggered at (1) the running onset or (2) 120 cm location during the cue-constant segment. The run onset stimulation is triggered at the first time point where the animals speed exceeded 10 cm/s for at least 200 msec after the trial start. The stimulation trial types were randomly selected for each stimulation session. The behavior analysis was performed using all the trials during the stimulation session. Only considering the stimulated trials led to similar results.

As a control, in some recording sessions, the optogenetics patch cables were only loosely mated with the implanted fiber optic cannulas, preventing light from propagating into CA1.

#### Acute extracellular electrophysiology

Two days before the recording, the animal was acclimated to the recording condition: including turning on the microscope light, removing Kwiksil that covered the skull, and adding saline to keep the craniotomy wet. On the day before the recording, craniotomy was opened. During electrophysiological recordings, a 64 channel silicon probe (neuronexus, buzsaki64sp) was slowly lowered into the hippocampal CA1 region. Data from all channels were filtered (0.3 Hz to 10 kHz), amplified (gain = 400) and continuously sampled at 20 kHz using the Amplipex system (16-bit resolution)^45^. Time stamps of behavioral events, and electrophysiological recording data were synchronized, recorded and stored on a computer.

For extracellular recordings with optogenetics, a 64 channel silicon probe with fibers mounted on the shanks (neuronexus, buzsaki64sp-OA64LP) was used. Photostimulation followed the same protocol as described in the optogenetics section above. The light source was a self-constructed laser diode array (6 diodes, 450 nm, osram-os, PL450B). Each diode was coupled to one shank on the probe. 2 to 5 diodes were coactivated during the stimulation. The stimulation laser power was < 1.2 mW at the tip of the probe.

For optogenetic tagging experiments, interneurons were tagged using 1ms pulses of blue light. Spike times around each pulse were binned with a resolution of 1ms and the significance of light responses in a 5ms window after the light pulse was assessed using the stimulus-associated spike latency test (SALT)^46^. Neurons with a P < 0.01 that responded to at least 40% of light pulses were identified as PV-positive. 30-40 tagging pulses were applied per recording.

#### Muscimol infusion

Muscimol hydrobromide (Tocris-0289) was dissolved in 0.9% saline. The control solution was saline. The muscimol injections were carried out as follows. First, an injection cannula was connected to a 10-μl Hamilton syringe through Tygon tubing (Tygon 720993) and filled with muscimol (1 mg ml−1) or saline (0.9%). Then the syringe was mounted into a microinjection pump (UMP3 with SYS-Micro4 controller). At the beginning of the injection procedure, the dummy cannula was removed from the guide cannula and replaced by the injection cannula (33 Gauge) which extended 0.5 mm deeper into the brain than the guide and dummy cannulae. Then 300-500 nl of muscimol or saline was slowly injected (100 nl/min) into CA1 and the injection cannula was left in place for another 3 min after the infusion was complete. Next, the dummy cannula that was cleaned with alcohol and dipped in sterile mineral oil was inserted back into the guide cannula. During the infusion, animals performed the PI task with water automatically delivered at the end of each trial. Animals did not show signs of stress or discomfort during the procedure. The animals were released from the setup after the infusion, and ∼30 min after the completion of the infusion, animals were placed back onto the VR setup. A typical muscimol infusion recording was composed of one control session before the infusion, and multiple recording sessions at 30 min, 1 hour, 2 hours after the infusion. Sessions after infusion were combined as “Musc” sessions in the analysis.

#### Spike sorting

To identify spikes, we performed off-line spike sorting on the recorded files following published methods using Klusters^47^ and Kilosort^48^. Units were further selected based on the percentage of spikes that violated refractory period and the mahalanobis distance from other units.

#### Histology

Mice were perfused transcardially with PBS followed by 4% PFA. Brains were post fixed overnight and transferred to PBS before sectioning using a vibratome. Coronal 50 μm free-floating sections were processed using standard fluorescent immunohistochemical techniques. PV immunoreactivity (Swant, GP72; 1:2000, guinea pig). The primary antibody was visualized by Alexa Fluor 488 AffiniPure Donkey Anti-Guinea Pig IgG (H+L) (1:1,000, Jackson ImmunoResearch Laboratories, 706-545-148). SST immunoreactivity (Millipore, MAB354; 1:250, rat). The primary antibody was visualized by Alexa Fluor 488 AffiniPure Donkey Anti-Rat IgG (H+L) (1:1,000, Jackson ImmunoResearch Laboratories, 712-545-153).

### QUANTIFICATION AND STATISTICAL ANALYSIS

#### Behavior analysis

To calculate how the speed changed over distance during the cue-constant segment, the histogram of speed within each trial was calculated using 1 mm distance bins. 15 trials were randomly selected from each session, and the average and SEM of speed were calculated across all the selected trials from all the sessions in each task. Similarly, to calculate how the number of licks changed over distance, a lick histogram with a bin size of 1 cm was first calculated for each trial. Then the same calculation as the speed was applied to get the average and SEM. To calculate the overall mean speed or lick within a certain distance range, the mean speed or lick number was first calculated for each session, then averaged across sessions from the same task. To calculate total stop time, we first smoothed running speed using a moving average with an 80ms window, and then summed the time when the speed was lower than 1 cm/s from the running onset to the end of the trial.

We found recordings with matching running speed between the passive and PI tasks, and identified 4 animals (10 recordings in total) where the mean speed did not differ from the passive task recordings.

#### Extracellular recording analysis

##### Alignment with sensory or motor cues

For both the passive and PI tasks, both the neuronal activity and the behavior of each trial were aligned with either the start cue onset or the running onset. The running onset for each trial was defined as the onset of the first running bout that lasted more than 0.3 sec with a speed > 10cm/s, starting from the previous trial after the animal had traveled the 180 cm cue-constant segment. The onset was when the running speed reached 1 cm/s before the running bout and after 180 cm in the cue-constant segment of the previous trial. If the speed never reached 1 cm/s, the time point when the speed was the lowest was considered as the onset, and this trial was classified as “no clear run onset”.

##### Firing rate profile of each neuron

Pyramidal neurons were identified based on their spike waveforms, and if their firing rates fell between [0.15 7] Hz. The spike train of each pyramidal neuron was first smoothed with a Gaussian function (s.d. = 30 ms) and then averaged across trials to get the average firing rate profile of that neuron. When examining how the firing rate changes over distance, the spike train was first binned using 1 mm spatial bins, and then smoothed with a Gaussian function (s.d. = 20mm). Normalized firing rate profiles of PyrUp or PyrDown neurons were calculated by first normalizing the firing rate profile of each neuron by its max firing rate and then averaging across the selected population.

##### Firing field identification

The averaged firing rate profile of each neuron was used to identify firing fields (internally generated fields). To identify firing fields over distance, the following criteria were applied: 1) a minimum mean firing rate of 0.09 Hz; 2) minimum mean trial-by-trial Spearman correlation of the firing rate profile > 0.15 and minimum spatial information^49^ > 0.25 bits/s; or minimum mean trial by trial correlation > 0.09 and minimum spatial information > 0.7 bits/s. 3) total number of trials > 15, and the neuron was active in at least 40% of the trials. 4) the maximum field width was 150 cm, measured by the width when the firing rate fell below 10% on both sides of the peak firing rate. 5) there was no other peak within 30 cm outside the field. The parameters were chosen based on visual inspection.

To identify firing fields over time, we used similar criteria as mentioned above. Some of the parameters were readjusted based on visual inspection: 1) minimum mean trial-by-trial correlation > 0.12 and minimum temporal information > 1 bits/s; or minimum mean trial-by-trial correlation > 0.09 and minimum temporal information > 2.5 bits/s. 2) The maximum field width was set to 4.5 sec. 3) There was no other peak within 1.7 sec outside the field. For the immobile task, the same method was used to identify firing fields over time with slightly different parameters. Neuronal activity was aligned with the last lick of each trial in the immobile task, and the following parameters were used: minimum mean trial by trial correlation > 0.1 and minimum temporal information > 0.65 bits/s.

##### Auto-correlogram

The spike time auto-correlogram of each neuron was calculated after binning its spike train using 1ms time bins.

##### Phase precession

Phase precession was calculated as the difference between the theta modulation frequency of individual neurons and the theta frequency of the local field potential (LFP). The theta modulation frequency of individual neurons was calculated using the peak time of its auto-correlogram within 100-200 ms and −200--100 ms windows^50^. One LFP trace recorded from the CA1 pyramidal layer was filtered with a third order Chebyshev Type II filter (4-16 Hz). LFP instantaneous frequency in the theta band was calculated using the Hilbert transform. The instantaneous frequency was then averaged over each trial and across trials to compute the LFP theta frequency.

##### Criteria for good and bad trials

The criteria for identifying good trials in the PI task are: 1) The animal came to a complete stop at the reward location of the previous trial just before the run onset (speed < 1 cm/s). 2) The subsequent running was mostly continuous, with a total stop time shorter than 2s after run onset. 3) The animal obtained a reward. Otherwise, the trial was considered a bad trial.

For the immobile task, trials where mice missed the reward or began licking before the halfway point of the delay period were labeled as bad trials.

##### Identification of PyrUp and PyrDown neurons

The change in firing rate around the run onset for each neuron was defined as the ratio between the averaged firing rate in a window 0.5 to 1.5 seconds (FRaft) and −1.5 to −0.5 seconds (FRbef) about the running onset (FRaft/FRbef). The PyrUp neurons were neurons whose FRaft/FRbef > 3/2, while the PyrDown neurons were neurons whose FRaft/FRbef < 2/3. PyrOther neuron were neurons whose FRaft/FRbef were between 3/2 and 2/3. The relative firing rate change of the average firing rate profile around run bouts and trial run onsets were calculated for each neuron in a window 0.5 to 1.5 and −1.5 to −0.5 seconds about the running onset with the formula (FRbef – FRaft)/(FRbef + FRaft).

The same criteria were used to identify PV-Up and SST-Up neurons.

To avoid bias in grouping neurons, an additional shuffling method was used to identify PyrUp and PyrDown neurons. Shuffled distributions of FRaft/FRbef were created by performing randomized circular shifts to the firing rate profile of each trial and recalculating FRaft/FRbef 1000 times. Neurons with a firing rate change that was above the 95th percentile or below the 5th percentile of the shuffled distribution were labeled as a PyrUp or PyrDown neuron, respectively.

The same method was used to identify PyrUp and PyrDown neurons with respect to the run onset in the passive task, and to the last lick in the immobile task.

##### PyrUp and PyrDown neurons with respect to running bouts in the PI task

Run bouts were identified as periods where locomotion exceeded 10cm/s for a duration of at least 0.5s. The onset of each run bout was traced back to the point where speed first exceeded 2 cm/s. If running speed in the window 0.5s before a run bout exceeded 10cm/s at any time point that run bout was excluded from analysis. Run bouts that occurred within the first 50 cm of a trial were also excluded from the analysis. Spike times for each neuron were aligned with the run bouts and firing rates were calculated in 2.5 ms bins and convolved with a Gaussian filter with a standard deviation of 30 ms.

Speed matching was performed for each recording by first calculating the mean speed and standard deviation of running bouts in a period from −1.5 to 0s and 0 to 1.5s around bout onsets. Trial run onsets that fell within one standard deviation of the mean speed for both periods were included for analysis. Recordings with less than 15 total run bouts and less than 15 speed-matched trial run onsets were excluded from the analysis.

##### Decay/rise time constant of PyrUp/PyrDown response

For each PyrUp neuron, the first peak in its mean firing rate profile that exceeded the 99 percentile of the shuffled mean firing rate profile (shuffled 500 times) was identified. The decay time constant was extracted by fitting the post-peak mean firing rate profile with the function a*exp(bx), where the time constant is defined as −1/b.

Similarly, for each PyrDown neuron, the significant peak 1 sec after run onset was identified as for PyrUp neurons. The time constant was calculated by fitting the function a*exp(bx) from the peak backwards to 1 sec after run onset.

##### Theta phase

The peaks, valleys, and zero crossing points of LFP were detected and assigned as phases 0, 180, and 90 or 270, respectively. Phases between these points were linearly interpolated^51^. Peaks in the LFP corresponded to the minimum firing of CA1 pyramidal cells. Please refer to Wang et.al. 2015^7^ for more details. Theta phase histogram of each neuron is calculated using 5-degree bins between 0 and 360 degrees.

##### Interneuron clustering

Interneurons were identified based on their spike waveforms and their auto-correlograms. Principle component analysis (PCA) was performed on the matrix formed by the normalized theta phase histograms of all the interneurons. The number of PCA components was selected such that the accumulated explained variance reached 95% unless the number of components was larger than 30. The same treatment was done on the matrix formed by the auto-correlograms of all the interneurons. The PCA components of the theta phase histograms, the auto-correlograms, and the estimation of the depth of interneurons relative to the center of the pyramidal layer were features used to perform kmean clustering of interneurons. The number of clusters was determined by the evalclusters function in matlab, and then went through visual inspection. The putative PV basket cell cluster was identified based on similarities with well-established firing phenotypes of this cell group^12,18^. The estimation of depths of interneurons and pyramidal neurons relative to the center of the pyramidal layer was calculated following the method in Mizuseki et.al., 2011^23^.

##### Decoding analysis

For each recording session, we trained a naïve Bayesian classifier built around the Matlab function fitcnb^28^ with population activity of all the PyrDown neurons without firing fields. Recordings with at least 30 good trials and 20 neurons that were either PyrUp or PyrDown neurons were included in the analysis. A uniform prior probability distribution was used. The PyrDown neurons were detected by thresholding FRaft/FRbef. Bayesian decoding posterior probability and decoding error were determined using tenfold cross-validation. Chance was determined by calculating the decoding error after performing randomized circular shifts of the firing rate profiles to each trial for N = 50 times. The time bin for decoding was 0.1 second.

##### Spectrograms and theta state changes

Multitaper spectrograms were constructed from the LFP channels in the layer center of CA1 using the mtspecgramc function in the Chronux library (http://chronux.org/). A sliding window of T = 1s and a frequency resolution of Δf = 2 Hz was used. The time-half-bandwidth product and the number of tapers used were calculated as in Prerau et.al. 2017^52^. Theta state was quantified by a theta/delta ratio, calculated as the quotient of the average power in the 6-8 Hz and 2-4 Hz bands.

##### Long pauses and inter-trial intervals

Long pauses were defined by immobile periods when running speeds were below 2cm/s for 5s or longer. For the purposes of brain state change analysis, running bouts were defined by periods when running speeds were 10cm/s or greater for a minimum of 3s. Inter-trial intervals were defined as the periods in between the delivery of reward in one trial and the running onset of the following trial. Slow runs were defined as periods where running speeds fell between 5 and 20 cm/s.

#### Statistics

No statistical methods were used to predetermine sample sizes, but our sample sizes are similar to those reported in previous publications. Data collection was not performed blindly to the conditions of the experiments, however the data collection and data analysis was performed by two experimentalists to ensure reproducibility of the results. For optogenetic experiments, stimulation types were randomly determined for each session. We used non-parametric statistical tests to estimate significant differences between groups: Wilcoxon rank-sum test and Kruskal-Wallis test. The Wilcoxon rank-sum test is two sided. All shuffling was done over 1,000 iterations.

